# Dysregulated circRNA expression profile and associated miRNA sponging in abnormal lung development in congenital diaphragmatic hernia

**DOI:** 10.1101/2025.10.13.681804

**Authors:** Marietta Jank, Arzu Ozturk Aptekmann, Muntahi Mourin, Nolan De Leon, Matthew Kraljevic, Daywin Patel, Wai Hei Tse, Claire McCallum, Richard Wagner, Shana Kahnamoui, Yuichiro Miyake, Athanasios Zovoilis, Michael Boettcher, Richard LeDuc, Richard Keijzer

## Abstract

**Rationale:** Circular RNAs (circRNAs) can function as disease biomarkers. Their profile in abnormal lung development in congenital diaphragmatic hernia (CDH) is unknown.

**Objective:** To evaluate circRNA expression profile in CDH-associated abnormal lung development.

**Methods:** We profiled circRNAs in rat CDH and control lungs at embryonic day (E)15 and E21 by microarray. We validated identified circRNAs using back-splice junction amplicon sequencing, RT-qPCR, and *in situ* hybridization. We modified a *CircRNA Function prediction Tool* to predict CircRNA::micro(mi)RNA::messenger(m)RNA interactions and compared these with Oxford Nanopore RNA sequencing and existing human CDH datasets.

**Measurements and Main Results:** Microarrays revealed a unique circRNA biosignature during CDH lung development. CircAnp32e was expressed in a sex-specific and spatiotemporal expression pattern in the epithelium at E15. The predicted mature sequence of circAnp32e overlapped >90% with its human orthologue. CircRNA::miRNA::mRNA interaction networks in E15 and E21 revealed enrichment in inflammation/infection, smooth muscle cell function, cell proliferation/cell cycle regulation, and response to hypoxia pathways. Parental genes of differential expressed circRNAs at E15 enriched pathways linked to cell proliferation/cell cycle/cancer, while at end-gestation, inflammation and cardiovascular processes were also overrepresented. Rat and human CDH lungs showed overlapping pathways with additional enrichment for RNA processing and protein binding/modification in humans.

**Conclusion:** A unique circRNA signature during abnormal lung development in CDH may mediate inflammatory responses, smooth-muscle-cell function, and cell proliferation regulation via miRNA sponging. Overlap of downstream pathways in rat and human CDH suggest conserved functions across species. CircRNAs may serve as biomarkers to guide prenatal management and mitigate aberrant lung development in CDH.

## Introduction

Lung development requires a delicate balance of cell proliferation, migration, and differentiation.^1^ This balance is disturbed in congenital diaphragmatic hernia (CDH) causing cardiopulmonary hypoplasia, pulmonary hypertension and a diaphragmatic defect.^2^

Circular RNAs (circRNAs) regulate gene expression and cellular processes in development and disease.^3,4^ CircRNAs have a covalently closed-loop configuration generated by backsplicing, which provides them with remarkable stability.^3^ circRNAs can serve as micro(mi)RNA sponges, protein scaffolds, and peptide templates via non-canonical translation, while modulating transcription, alternative splicing, and mRNA translation to regulate cell proliferation, apoptosis, and stress responses.^3^

CircRNAs show a tissue- and developmental stage-specific expression suggesting they can serve as ideal biomarkers for congenital anomalies. We previously published a unique circRNA biosignature in the lungs of deceased CDH cases at mid- and end-gestation.^5^ We have published dysregulation of a circRNA derived from the parental gene *Tial1* in the nitrofen rat model for CDH at late gestation and established a modified bioinformatics pipeline to predict mature circRNA sequences and biological functions through miRNA sponging.^6^ Here, we aimed to elucidate and validate circRNA expression profiles across abnormal lung development in CDH and to infer their potential biological functions.

## Methods

### Samples

Procedures were approved by the Animal Research Review Committee (23-026 [AC11831]) and human Research Ethics Board (HS15293 [H2012:134]) at the University of Manitoba. The nitrofen model was employed as previously described.^6^ Lungs were harvested at E15 and at E21 only from fetuses with confirmed diaphragmatic defects. Sex was determined using X- and Y-specific primers (Table E1).

### RNA isolation, RNase R treatment, and DNA isolation

Total RNA was extracted from fetal lungs at E15 and E21, quantified, and enriched for circRNAs by RNase R digestion. Genomic DNA isolation was performed with in-column RNase A digestion. See supplement for details.

### Microarray expression of circRNAs

CircRNAs were labeled using random primers and hybridized to Arraystar Rat Circular RNA Microarrays (Ref. AS-S-CR-R-V2.0, Arraystar Inc.). See supplement for details.

### cDNA library generation, circRNA back-splice junction detection, subcloning, and Sanger sequencing

Primers were designed in-house, and PCR was performed on diluted RNase R-treated cDNA after cDNA synthesis. Plasmids from positive colonies were sequenced and analyzed once ligation products were transformed and selected. See supplement for details.

### RT-qPCR

Following primer optimization and sequence verification, RT-qPCR was performed on diluted cDNA (1:100 for parental genes and 1:10–1:20 for circRNAs). See supplement for details (Table E2).

### BaseScope™ in situ Hybridization

Sections were hybridized with BaseScope™ probes (Table E3) and amplified following a modified manufacturer’s protocol (Table E4).

### Oxford Nanopore Sequencing

RNA was extracted as described above and sequenced. For a detailed protocol of the Oxford Nanopore sequencing experiment, see Table E5 and supplement.

### Bioinformatics analysis and modified CRAFT

Downstream targets as predicted by our modified CRAFT pipeline were enriched for Gene Ontology (GO) and KEGG pathways.^7,8^ Pathways were compared with direct RNA sequencing and human CDH datasets.^5^ See supplement for details.

### Statistical Analysis

We quantified in situ hybridization signal per cell in four random areas with HALO® (Indica Labs, Albuquerque, NM, USA). Statistical analyses and data visualization were performed using GraphPad Prism version 9.0.2 (GraphPad Software, San Diego, CA, USA) and R version 4.3.2 (R Core Team, 2023).

## Results

### CircRNA expression profile at E15 and E21

Microarray analysis of rat fetal left lungs detected 13,337 and 13,770 circRNAs at E15 and E21, respectively. Principal component analysis (PCA) revealed clustering of nitrofen-induced and control lung datasets at both stages (E15: Figure E1A and E21: Figure 1B). At E15, differentiation was visible between CDH and control lungs within differentially expressed (DE) circRNAs (Figure E1B, E1C). At E21, this distinction extended to all circRNAs, regardless of significance (Figure 1C, 1D). At E15, 119 circRNAs were significantly DE, with 22 passing Benjamini-Hochberg adjustment. At E21, 681 were DE, and 163 remained significant after Benjamini-Hochberg adjustment. Full results for both embryonic stages are available upon request.

**Figure 1.**
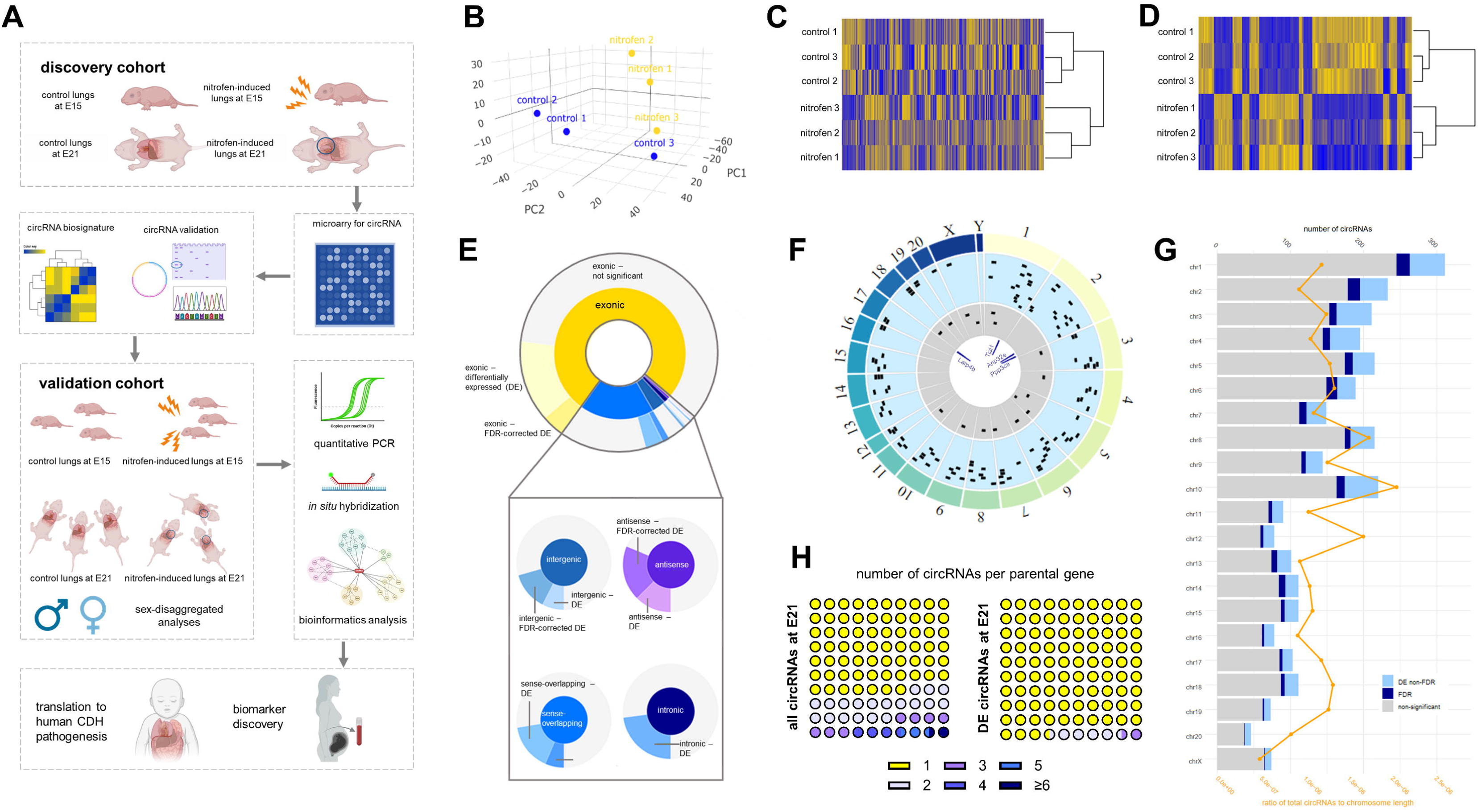
Unique biosignature for lung hypoplasia in CDH. Graphical abstract depicting the evaluation of the circular RNA (circRNA) profile in early and late stages of abnormal lung development in the nitrofen rat model for CDH using a microarray approach (A). Grouping of control and nitrofen-induced lung samples at embryonic day (E)21 based on overall circRNA profile plotted on a principal component analysis (B) and heatmap of all detected circRNAs (C). A unique circRNA biosignature of differentially expressed (DE) circRNAs distinguishes control and CDH lungs at late gestation (D). Distribution of the circRNA type at E21 where most of the DE circRNAs (71%) are exonic or sense-overlapping (22%) (E). Chromosomal location of the back-splice junctions at E15 (inner circle) and E21 (outer circle) showing labeling of the circRNA candidates chosen for validation (F). The number of circRNAs across all chromosomes and the ratio of circRNAs relative to the chromosome length, indicating highest enrichment of chromosomes 10, 8, and 12 at E21 (G). Number of circRNAs per parental gene for all circRNAs and DE circRNAs at late gestation (H).

Exonic circRNAs accounted for 77% and 71% of DE circRNAs at E15 and E21, respectively. Sense-overlapping circRNAs comprised 23% (E15) and 22% (E21), followed at E21 by intergenic (5%) and antisense (2%) circRNAs (Figures E1D, 1E). To identify chromosomal “hotspots,” we mapped circRNA coordinates and normalized by chromosome length. Similar patterns were evident at E15 (Figure E1E) and E21 (Figure 1G). Chromosome 1 showed the highest circRNA count at E21, while chromosomes 10, 8, and 12 had higher relative densities. Chromosome X had the lowest ratio (4.85e^-07^). Multiple circRNAs originated from single parental genes (Figures 1H, E1F). At E21, dcc produced 12 isoforms. Among the DE circRNAs, 9 had 2 isoforms from the same parental gene (*dcc, ddhd2, ezh2, nckap5, ptk2, rn18s, stk39, trps* and *wdr7*).

### Validation of candidate circRNAs from parental genes anp32e, larp4b and ppp3ca using subcloning and directional PCR

Based on microarray findings and relevant literature, we validated circRNAs derived from the parental genes *anp32e*, *ppp3ca*, *larp4b* and *tial1*^6^ (Table 1).

**Table 1.**
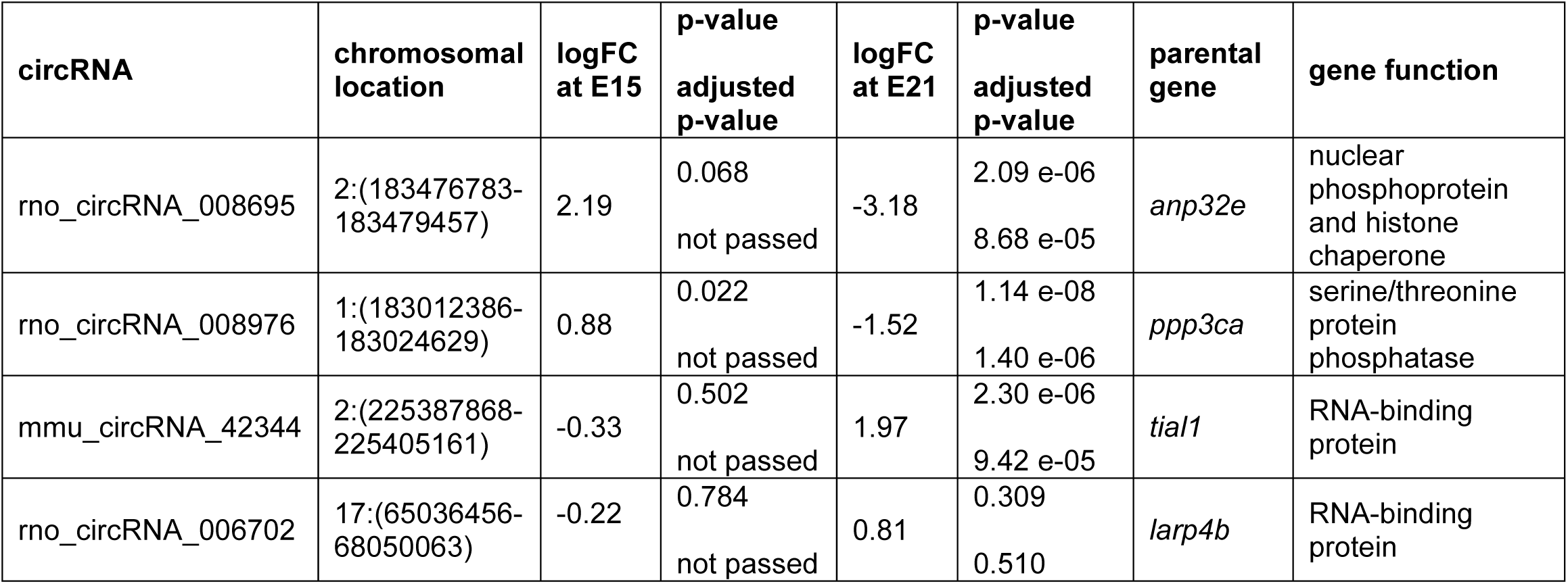
Selected circRNA candidates.

circAnp32e had the lowest logFC at E21 (-3.18) and is linked to fibrosis, epithelial– mesenchymal transition (EMT), and viral life cycles.^9,10^ circAnp32e spans exons 4-2 on rat chromosome 2 and its back-splice junction was confirmed using in-house designed divergent primers on RNase R-treated cDNA, followed by subcloning and Sanger sequencing (Figure 2B). Considering circRNAs are generated through post-transcriptional backsplicing events and therefore absent in linear genomic DNA, back-splice junction was only amplified in the cDNA pool with and without RNase R treatment (Figure 2B). We established directional PCR amplification using divergent primers validating alignment of exon 4-3-2 with head-to-tail splicing sites by Sanger sequencing (Figure 2E). This matches the 442-nucleotide mature sequence of circAnp32e(2,3,4).1 annotated in the circAtlas database (rno-Anp32e_0001; Figure E2A accessed 28.7.2024).

**Figure 2.**
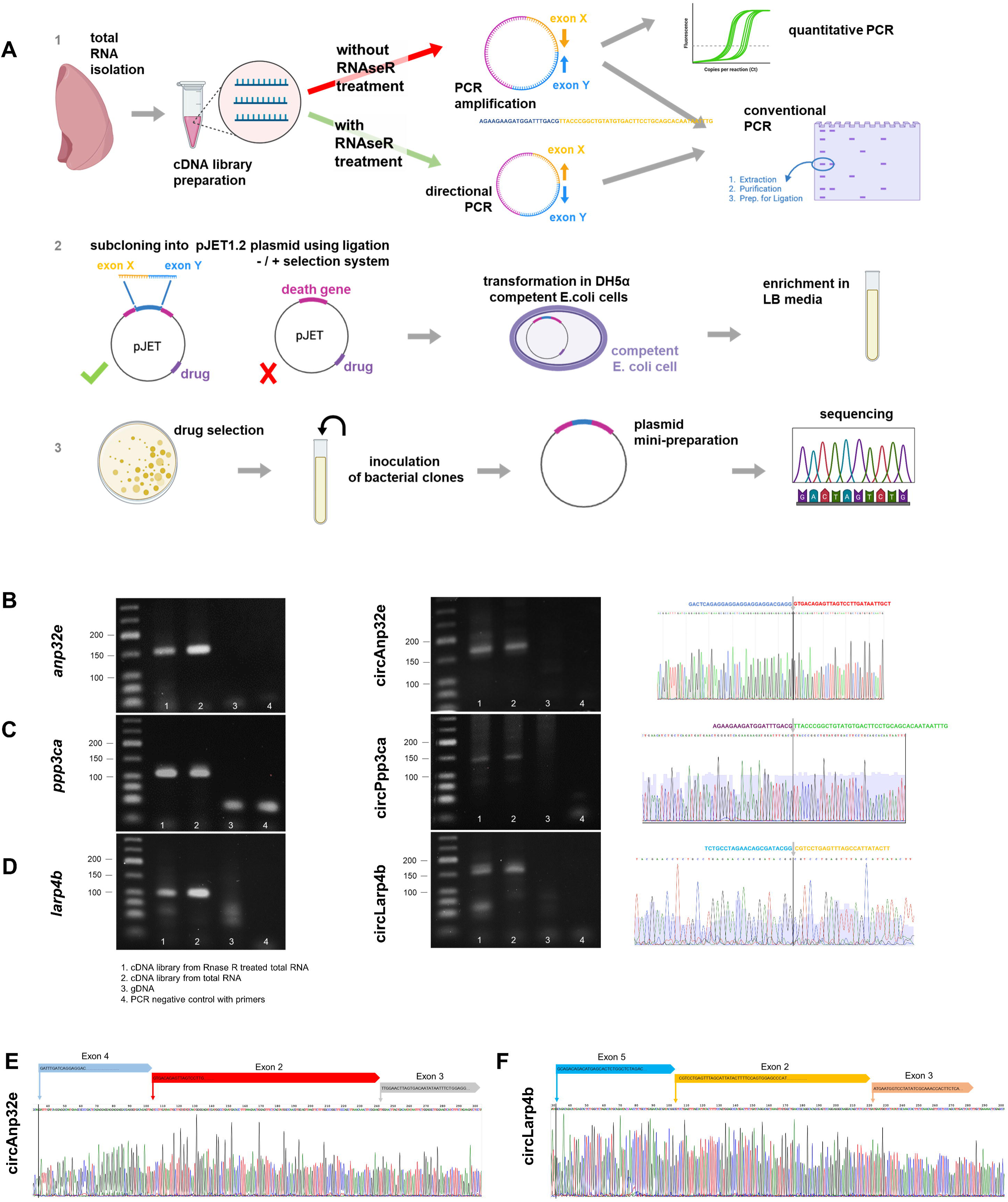
Validation of circAnp32e, circPpp3ca and circLarp4b by conventional PCR with convergent and divergent primers using pJET subcloning and amplicon sequencing. Total RNA was isolated and reverse-transcribed; cDNA was treated with or without RNase R. Untreated samples enabled quantification by qPCR, while RNase R treatment depleted linear RNAs facilitating junction-specific PCR and gel electrophoresis (A–1). PCR products were subcloned into the pJET1.2 vector, transformed into competent cells and enriched in selective media (A–2). Clones with inserts were selected via antibiotic resistance, followed by plasmid mini-preparation and Sanger sequencing for validation (A–3). This workflow was applied to candidate circRNAs and their parental genes using different experimental conditions: (1) cDNA synthesized from RNase R-treated total RNA to selectively enrich for circRNAs, (2) cDNA synthesized from total RNA to account for linear RNA amplification and normalization to a housekeeping gene for quantification by RT-qPCR, (3) genomic DNA (gDNA) as a control for potential DNA contamination, and (4) PCR negative control without a template to assess non-specific amplification. Divergent primers successfully amplified the circRNA derived from the parental genes *anp32e, ppp3ca* and *larp4b* in RNase R-treated cDNA but not in genomic DNA (B, C and D, respectively). Directional PCR amplification detected the additional circRNA exonic junctions for circAnp32e and circLarp4b to prove the circular nature of the RNA molecule in rat lung tissues (E and F, respectively).

circPpp3ca was DE at E21, and aberrations in its parental gene have been linked to multiple congenital abnormalities.^11^ Its back-splice junction composed of exons 10-7 on rat chromosome 2 was confirmed (Figure 2C; the form is also annotated in circAtlas Figure E2B). Similarly, we validated circRNA derived from *Larp4b*, an alternative splicing-related RNA-binding protein. circLarp4 expression correlates with non-small cell lung cancer prognosis and suppresses cancer cell proliferation.^12,13^ Sanger sequencing confirmed the back-splice junction between exons 5 and 2 (Figure 2D) and exon alignment 5-2-3 (Figure 1F).

### circRNA and parental gene expression profiles and spatiotemporal distribution of candidate circRNAs: circAnp32e, circLarp4b, and circPpp3ca

As circRNAs may co-regulate their linear host transcripts^3^, we quantified both parental genes and circRNAs from the same cDNA pool via RT-qPCR (Figure 3A and 3B). *Anp32e* was downregulated in nitrofen lungs at E15 and E21; the interaction factor for sex in two-way ANOVA was not significant (p=0.75). RNA sequencing confirmed *anp32e* downregulation without reaching statistical significance (logFC = -3.5, p=0.15; Figure E3A). At E15, overall circAnp32e expression was similar between controls and CDH samples (p=0.22); however, expression was upregulated in male lungs (p=0.034, Fig. 3B). Consistent with our microarray studies, RT-qPCR revealed a decreased circAnp32e expression in nitrofen-induced lungs at E21 (p=0.004). *Ppp3ca* was upregulated at E21, an effect driven by expression in male lungs (p=0.006; Figure 3A). Consistent with microarray results, nitrofen lungs had decreased circPpp3ca expression at E21 (p=0.016). Overall *Larp4b* and circLarp4b expression did not differ between control and nitrofen-induced lungs, except for significantly elevated levels in male CDH lungs based on a sex-disaggregated analysis (in line with the microarray data).

**Figure 3.**
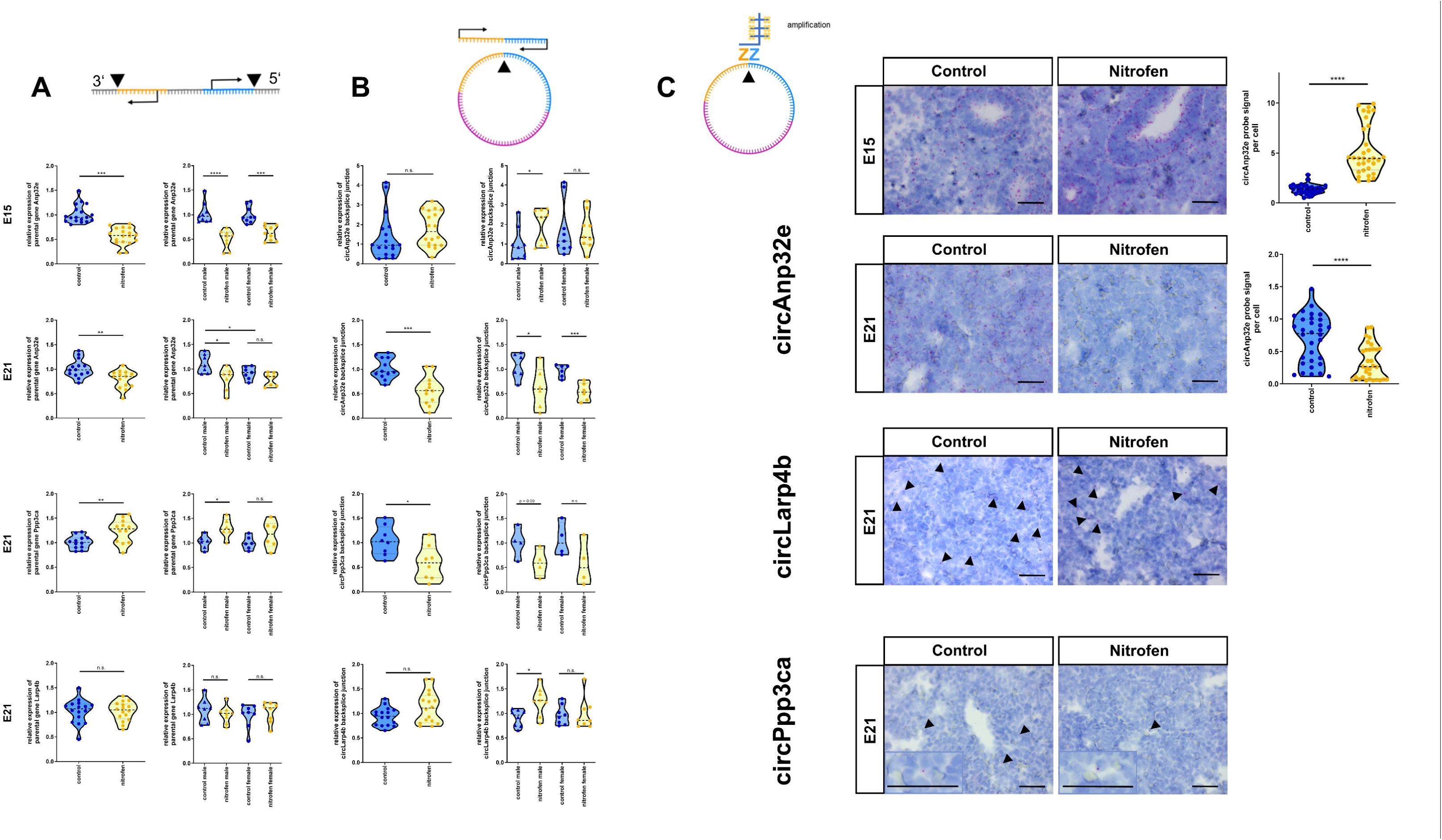
Sex- and developmental stage-specific expression of selected circRNA candidates. Linear mRNA transcripts are often co-expressed with corresponding circRNAs, which in turn can regulate the expression of parental genes. Therefore, we performed quantitative PCR of both the parental gene (A) and circRNA (B) for the following candidates (top to bottom): (circ)Anp32e (E15), (circ)Anp32e (E21), (circ)Ppp3ca (E21) and (circ)Larp4b (E21). To refute a potential overestimation or false-positive detection resulting from PCR-based methods, we further confirmed the microarray results using *in situ* hybridization (C). *\*P*<0.05, *\*\*P*<0.01, *\*\*\*P*<0.001.

*In situ* hybridization confirmed circAnp32e and circLarp4b expression (Figure 3C). The circAnp32e signal peaked in the epithelium of nitrofen lungs at E15. CircPpp3ca showed a specific but weak probe signal in both groups corroborating the high Cq values in RT-qPCR; quantifying specific spatiotemporal patterns was not feasible (Figure 3C).

### Bioinformatics analyses of predicted circRNA function through miRNA sponging in abnormal lung development

To explore circRNA function in abnormal lung development, we performed GO and KEGG pathway analyses of parental genes and circRNA::miRNA::mRNA targets at E15 and E21.

At both E15 and E21, 730 circRNA::miRNA::mRNA target genes enriched for various pathways, with complete overlap among the top 15 KEGG terms (Figure 4B). Additional terms related to vascular development and endothelial migration (biological processes), NF-κB signaling (molecular function), and membrane and focal adhesion structures (cellular components) enriched at E15 (Figure E4B). At late gestation, 583 predicted downstream mRNA targets augmented biological processes linked to cellular response to hypoxia, regulatory networks in smooth muscle cell proliferation and cardiac muscle development, as well as apoptosis signaling and interleukin-1 response. Enriched molecular functions included cytokine receptor binding and activity, cellular adhesion, and extracellular matrix function, while enriched cellular components comprised collagen matrix, muscle fiber components, and membrane structure (Figure 4A).

**Figure 4.**
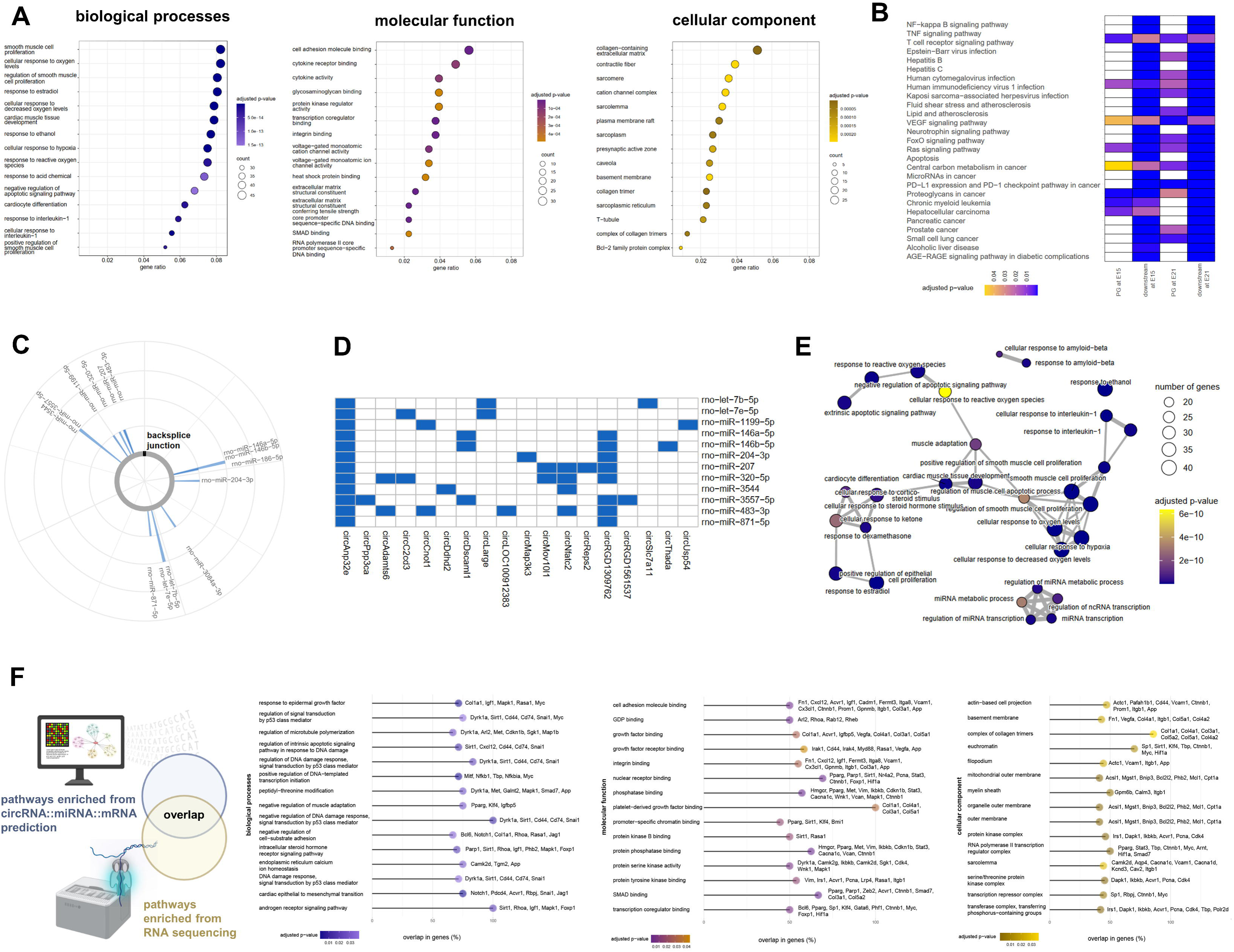
Enrichment for response to hypoxia, smooth muscle cell proliferation, and inflammation. Enrichment of top 15 gene ontology (GO) terms from circRNA::miRNA::mRNA downstream targets at E21 focuses on muscle cells, extracellular matrix, response to hypoxia, and binding (A). The heatmap compares the top 15 KEGG pathways enriched by parental genes and circRNA::miRNA::mRNA targets at E15 and E21, suggesting a substantial overlap in predicted biological functions by miRNA sponging at early and late gestation (B). Interaction of circAnp32e with miRNAs (C). Heatmap summarizing circRNAs interacting with the same miRNAs as circAnp32e (D). Pathway analyses of mRNA targeted by this circRNA network (E). Oxford Nanopore sequencing was utilized to confirm predicted pathways and identify promising downstream targets. The plots show the overlap in significantly enriched GO terms, the proportion of DE genes identified through RNA sequencing and their respective names (F).

The parental genes of DE circRNAs at E15 enriched for splicing/mRNA processing, and (mesenchymal/stem/neural crest) cell development as well as migration (Figure E4A). Enriched GO terms for molecular functions were predominantly binding-related, while cellular components included RNA-regulatory complexes (Figure E4A). While at E21, we found enriched GO terms related to muscle hypertrophy and a unique cluster around neuron development (Figure E4C). Both embryonic stages were associated with KEGG pathways related to inflammation and infection, as well as cancer, cell cycle regulation, and proliferation (Figure 4B).

Next, we analyzed miRNA sponging by circAnp32e (Figure 4C) and DE circRNAs with shared miRNAs (Figure 4D). CircAnp32e interacted with miRNAs known to influence lung development in CDH, including miRNA-let-7b-5p and miRNA-let-7e-5p (Figure 4C).^14^ The heatmap summarizes circRNAs that interact with the same miRNAs as circAnp32e; one of which is circPpp3ca (Figure 4D). Pathway analyses of mRNA targeted by this circRNA network revealed GO terms associated with smooth muscle and epithelial cell proliferation, apoptosis, and miRNA transcription (Figure 4E).

Finally, validation using Oxford Nanopore RNA sequencing confirmed overlap in enriched GO terms between the predicted circRNA::miRNA::mRNA targets and observed expression changes (Figure 4F and Figure E7A). Not only did the enriched biological processes linked to apoptosis, epithelial-mesenchymal transition (EMT) and muscle adaptation overlap, genes contributing to these pathways showed > 60% similarity (Figure 4F). All enriched KEGG terms from the CRAFT-based prediction and RNA sequencing presented a moderate positive correlation (Figure E7B).

### Indications of overlap in circRNA profile and function between human and rat CDH at later stages of abnormal lung development

Given the evolutionary conservation of circRNAs, we compared expression profiles between rat CDH lungs at E21 and our previously published human CDH lung profiles at mid- and end-gestation.^5^ Human CDH showed higher total circRNA enrichment for chromosomes 17 and 19 (Figures E5A–B) and DE circRNAs clustered on chromosome 1 (Figure 5C). Shared parental genes of DE circRNAs in rat and human CDH were associated with lung disease, embryonic development, fibrosis, proliferation, and RNA/DNA regulation (Table E6).

**Figure 5.**
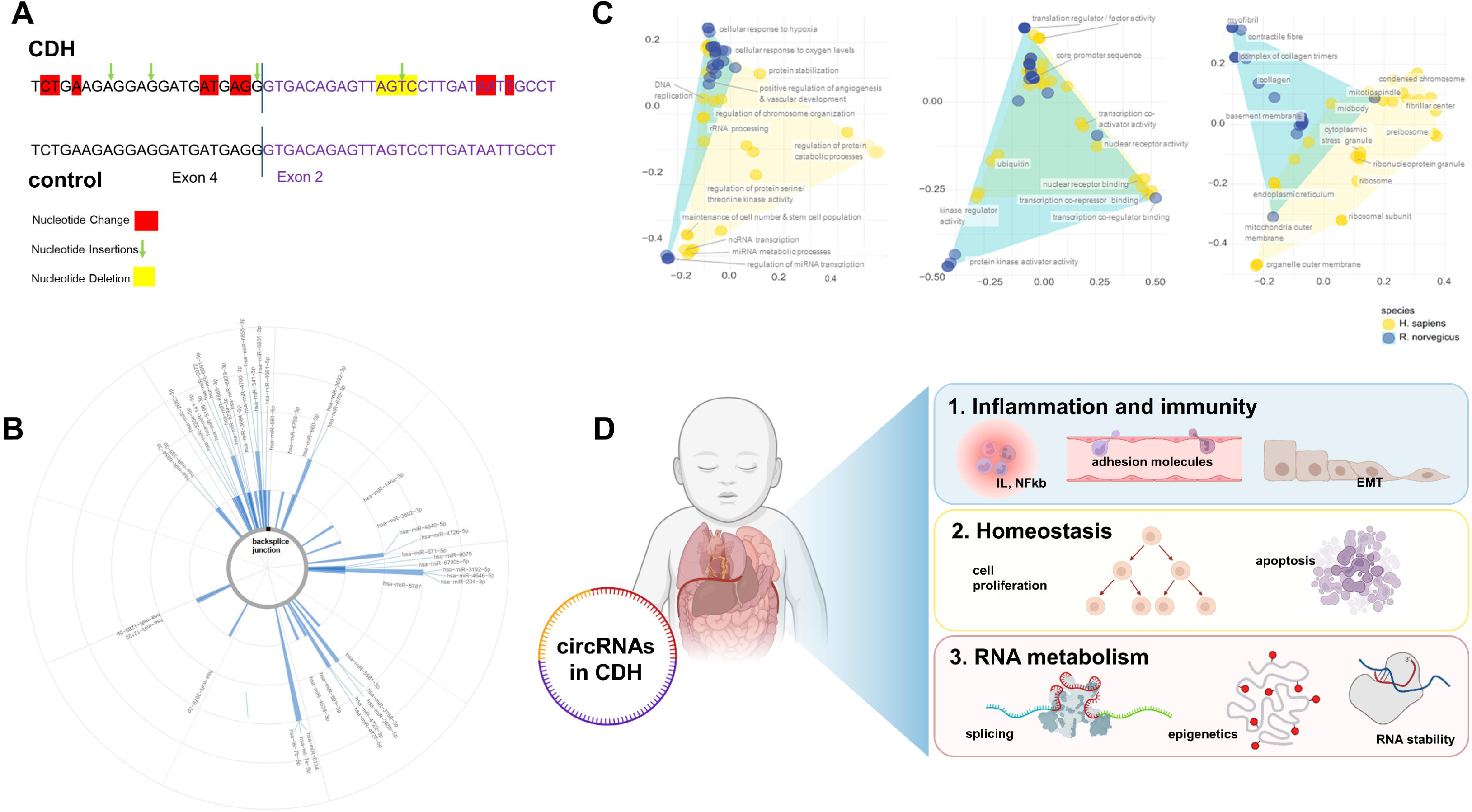
Conserved circRNA sequence and strong overlap in biological function (miRNA sponging) in rat and human CDH. We compared the circRNA profile in human CDH lungs at mid- and end-gestation with the findings in our nitrofen rat model. The circAnp32e mature sequence is highly conserved between the two species, and we successfully detected circAnp32e in formalin-fixed paraffin-embedded tissue of CDH lungs. Within the circAnp32e back-splice junction, the sequence detected in CDH lungs revealed more nucleotide variations compared to control samples (A). Further, human circAnp32e is predicted to sponge similar miRNAs to the rat circRNA equivalent shown in Figure 4C (B). There was a strong overlap for the gene ontology (GO) terms between the two species; in particular for binding, RNA processes, response to hypoxia, and vascular development (C). circRNAs may be the missing link in the pathogenesis of CDH connecting genetic susceptibility to environmental factors, and modulating pathways associated with inflammation/immunity, regulation of epithelial/mesenchymal/smooth muscle cell proliferation, and RNA processing by miRNA sponging (D).

Predicted circRNA::miRNA::mRNA targets demonstrated conserved enrichment in GO terms between the two species (Figure 5D), including (1) cellular response to hypoxia and regulation of miRNA transcription (biological processes), (2) transcription factor binding and regulation of translation activity (molecular functions), and (3) chromosome-associated components (cellular components). Shared KEGG terms enriched for inflammation and infection (*e.g*., NF-κB and TNF signaling, viral infections), regulation of cell cycle and proliferation, as well as cardiovascular pathways (Figure E6C). Interestingly, only human CDH showed enrichment for epigenetic pathways (Figure E6C).

Lastly, we evaluated the human orthologue of circAnp32e for comparative analysis. Rat circAnp32e(2,3,4).1 (or rno-Anp32e_0001) aligned 90% with human circANP32E(2,3,4).1 (hsa-ANP32E_0007) (Figure E5C). PCR amplification and Sanger sequencing successfully confirmed its expression in human fetal lungs (Figure 5A). Notably, we observed nucleotide variations within the disease-specific sequences that were absent in the control samples. Further, both orthologues are predicted to sponge similar miRNAs (Figure 5B, compared to 4C in the rat model).

## Discussion

The etiology of CDH remains unknown but presumably involves genetic susceptibility and environmental factors that may trigger epigenetic modifications affecting gene expression during critical developmental stages.^2^ Among these, DNA methylation^15^, histone acetylation^16^ and non-coding RNAs^5,17^ have been implicated in CDH. Here, we investigated the circRNA expression profile and function (miRNA sponging) in early and late abnormal fetal lung development. We identified sex-specific expression of circAnp32e, circPpp3ca, and circLarp4b, with spatiotemporal localization of circAnp32e in the pulmonary epithelium. We demonstrated for the first time conserved circRNA forms and functions in the nitrofen rat model and human CDH. Our findings provided novel insights into CDH pathogenesis and identified specific DE circRNAs as potential new CDH biomarkers and therapeutic targets.

Although circRNA expression has been characterized during physiological rodent lung development^22–24^, our study identifies a distinct circRNA biosignature in CDH-associated aberrant lung development. In line with reports that circRNAs are more abundant in fetal than adult tissues and exhibit strong tissue- and stage-specificity^18,19^, our data reveal pronounced developmental dynamics. At E15, we identified broader ranges of circRNA isoforms produced per parental gene suggesting that circRNAs may regulate fundamental processes such as cell proliferation, differentiation and lineage specification in early development. As development progresses, circRNA-mediated regulation appears to also impact more specialized functions related to tissue maturation and organogenesis, including responses to environmental factors (*e.g*., infection/inflammation, hypoxia) (Figure 4A-B). Nanopore sequencing revealed a 1.7-fold increase in total transcripts from E15 to E21, and an additional 1.3-fold increase after nitrofen exposure. Correspondingly, the number of DE circRNAs increased, although the circRNA isoform diversity decreased at E21 – suggesting intricate temporal regulation (Figure E1F). Similar trends have been observed during porcine brain development.^20^ These circRNA “hotspot” loci may reflect conserved regulatory elements that are modulated through cis-regulatory and epigenetic mechanisms, and their dysregulation could overlap with disease-specific regions^25^; warranting future studies (Figure 5C). Furthermore, they may reflect underlying post-transcriptional regulatory mechanisms complementary to intron/exon-specific (cis)-elements within the parental gene that facilitate circRNA formation by interacting with RNA-binding proteins (RBPs). As such, different isoforms from the same parental gene can exhibit distinct expression levels. For example, *Ppp3ca* yielded 20 circRNAs at E15 but only four at E21, of which one was significantly dysregulated in CDH lungs. We propose that RBPs provide selective circRNA stability over time, thereby allocating regulatory functions to fewer, more critical isoforms as differentiation progresses.

To date, few studies have investigated circRNAs in neonatal lung conditions like bronchopulmonary dysplasia and CDH.^6,26,27^ Our results indicate that most DE circRNAs were exonic corroborating previous studies. Accordingly, we investigated their function (by sequestering miRNAs) and downstream mRNA regulation.^18,28^ At both E15 and E21, circRNA targets enriched pathways linked to inflammation, epithelial/mesenchymal/smooth muscle cell proliferation and differentiation, as well as response to hypoxia – this was confirmed by RNA sequencing. Abnormal pulmonary remodeling secondary to CDH is presumably driven by impaired lung growth, altered vascularization, and cellular processes such as epithelial- (EMT) and endothelial-to-mesenchymal transition (EndoMT).^2^ EMT and EndoMT involve the transition of epithelial and endothelial cells, respectively, to more mesenchymal cell-like phenotypes (*e.g*., myofibroblasts and smooth muscle cells) under physiological and pathological conditions.^29^ In CDH, elevated inflammatory pathways may facilitate EMT and EndoMT resulting in pulmonary fibrosis and remodelling of pulmonary vessels.^2,30–32^ CircRNAs, depending on their targets, may modulate inflammation, apoptosis, fibroblast proliferation, and extracellular matrix deposition.^33^ For example, during TGF-β1–induced EMT, circRNAs are dynamically regulated by splicing factors like Quaking (QKI)^29,34^, which is dysregulated in CDH lungs as we previously published.^35^ Supporting these findings, our data demonstrated that circRNAs derived from known drivers of EMT and EndoMT - EZH2, Akt3, Clasp2, EP300, and PTK2^29,36,37^ - were differentially regulated at E21. Additionally, increased circAnp32e expression in the epithelium of nitrofen-induced lungs led us to speculate on a regulatory role for this specific circRNA in CDH-associated EMT.

CircRNA sequences and miRNA binding are highly conserved across species. Indeed, circAnp32e showed conserved sequence and predicted miRNA interactions between rats and humans (Figure 5A and 5B). Further, predicted circRNA::miRNA::mRNA interactions at E21 matched GO/KEGG terms, including hypoxia response, transcription/translation regulation, and inflammatory/infectious pathways, at human mid- and end-gestation, suggesting potential conserved circRNA-mediated regulatory mechanisms (Figure 5D, Figure E6A-C). Considering the enrichment of cardiovascular pathways and adhesion molecule-related terms at E21 as well as the similarities in molecular/signaling mechanisms between EMT and EndoMT, it is reasonable to propose circRNA involvement in EndoMT. Emerging data from brain endothelium in mice after stroke or inflammatory stimulation^38,39^ and human umbilical vein endothelial cells subjected to sera from patients with Kawasaki syndrome^40,41^ further support this notion by showing regulation of EndoMT by circRNAs through miRNA sponging, thereby regulating tight junctions, and the expression of interleukins and mesenchymal markers. Despite strong associations, further validation for a functional role of specific circRNAs in EMT and EndoMT is still required. Our data suggest circAnp32e as a candidate for future mechanistic studies.

Our validated candidates (circAnp32e, circPpp3ca, circLarp4b, and circTIAL1^6^) show differential expression patterns diverging from their linear mRNA counterparts, in line with previous human fetal brain and rat lung development studies.^22,42^ circRNAs can regulate parental gene expression by stabilizing mRNAs or enhancing transcription via RNA polymerase II.^28^ ANP32E, PPP3CA, TIAL1 and LARP4B are linked to transcriptional regulation, apoptosis and chromatin modification. ANP32E is a highly conserved histone chaperone protein that is enriched in endothelial cells of human cardiac tissue and implicated in the life cycle of various viruses.^10,43,44^ PPP3CA is a calcineurin subunit involved in cardiac development/pathological remodeling, T-cell activation, and cell cycle control/apoptosis.^11,45^ Lastly, LARP4b and circLarp4b are involved in mRNA metabolism and cancer pathogenesis by regulating cell proliferation, migration, cell cycle, and apoptosis^46,47^; mechanistically, circLarp4b decreases cell proliferation and migration by regulating expression of its parental gene.^48^ LARP4B and TIAL1 are RNA-binding proteins (RBPs) and part of a regulatory network with other RBPs that modulates mRNA turnover and translation.^6^ Some RBPs auto-regulate their own circRNA production by binding specific sequences in their pre-mRNA, thereby facilitating back-splicing. These circRNAs may serve regulatory roles (*e.g*., via miRNA sponging) or directly interact with mRNA UTR elements or RBPs.^26,49^ Our data showed opposite expression dynamics of the parental genes *Tial1* and *Larp4b* and their circRNAs, particularly in males, suggesting RBP autoregulation and/or coordination within a broader network of splicing-associated RBPs in CDH.

Others have reported enrichment in clusters associated with biological functions of the parental genes for organ-specific circRNAs.^18^ circRNAs are enriched in cardiac remodeling^50,51^, during stress responses^37^, and in the fibrotic lung.^33^ We observed clustering of circRNA::miRNA::mRNA interactions and associated parental genes in several biological processes, including mRNA processing, development, and enzyme/RNA binding at E15, and smooth muscle cell proliferation at E21. Additionally, circAnp32e shared common miRNAs with other circRNAs, including circPpp3ca, whose targets augmented apoptosis-related pathways, smooth muscle cell proliferation regulation, and miRNA transcription. Our results suggest coordinated regulatory networks formed by circRNAs, miRNAs, and mRNAs, that involve parental gene expression.

CDH is more prevalent in males and emerging data – including unpublished results from our group – suggest higher susceptibility of male CDH patients to certain long-term outcomes, including those affecting the respiratory system. Here, we show a sex-specific expression of individual circRNAs in CDH. Similar results have been reported in rat brain and lung development.^22^ While no DE circRNAs were identified on the Y chromosome in either rat or human CDH, our data showed enrichment of estradiol-responsive pathways among circRNA targets at E21, indicating a potential role for sex hormones. Accumulating evidence suggests sex-specific differences in organ structure, development, and function, which may be partially explained by variations in transcription factor binding, with 37% of all genes exhibiting sex differences in at least one tissue.^52,53^ Further, sex-specific regulation of RBP activity or upstream splicing mechanisms may influence circRNA biogenesis independently of parental gene expression. Supporting this, circRNAs have been shown to regulate miRNA biogenesis in a sex-specific manner in neonatal lung conditions such as BPD.^26^

While circRNA-based therapies have been explored in cancer^54^, prenatal application studies remain limited due to potential off-target effects and safety concerns. However, considering their function in various molecular and cellular processes, circRNAs can be envisioned as potential (prenatal) therapeutic targets. Their stability, developmental stage- and tissue-specific expression, and detectability in maternal biofluids highlight their promise as non-invasive prenatal biomarkers to support personalized counseling and clinical management.^5,6^

### Limitations

Microarrays are well-suited for high-throughput screening of circRNAs via back-splice junctions, particularly in low-input samples like formalin-fixed paraffin-embedded tissue or biofluids. However, because these analyses depend on the availability of pre-designed probes, the detection of novel or low-abundance circRNAs is limited. Further, our investigation of circRNA function relied on bioinformatics prediction, which is particularly challenging for species beyond human and mouse. Illumina RNA sequencing provides broader circRNA identification and parental gene expression data but is limited by low read coverage, short read lengths, accurate quantification due to low abundance, and intricacies of complex bioinformatics analyses.

CircRNA detection and quantification remain technically demanding due to relatively low expression levels and potentially high false-positive rates. Therefore, we performed targeted amplification using samples with and without RNase R treatment. RT-qPCR validation was limited by low expression of some circRNAs, particularly circPpp3ca. Similar validation challenges have been reported in normal rat lung development.^24^ *In situ* hybridization confirmed our qPCR-based detection but lacked sufficient signal for quantitative assessment in circPpp3ca and circLarp4b. Nonetheless, this technique holds great potential for the detection of abundant circRNAs (*e.g*., circAnp32e) enabling spatiotemporal analyses.

## Conclusion

We show distinct circRNA biosignatures for abnormal lung development in our model of CDH with suggested conserved elements across rat and human CDH. CircRNA-mediated miRNA sponging appears related to inflammation/immunity and regulation of epithelial/mesenchymal/smooth muscle cell proliferation. Our data imply that circRNAs could constitute a mechanistic link between genetic predisposition and environmental insults, and (pending further validation) may serve as prenatal biomarkers for timely detection of abnormal lung development.

## Acknowledgements

We thank the animal facility staff for their dedicated care and husbandry. The authors acknowledge Science Impact (Winnipeg, Canada) for (post-) editing of the final manuscript. Further, we thank Phil Snarr for supporting the submission process.

