## Supplementary material for "Dysregulated circRNA expression profile and associated miRNA sponging in abnormal lung development in congenital diaphragmatic hernia": Online Data Supplement

### Supplementary Methods

We investigated circular RNA (circRNA) dysregulation in the nitrofen rat model of congenital diaphragmatic hernia (CDH) at embryonic days (E)15 and E21 using an integrated experimental–computational workflow, followed by cross-species comparison with human CDH. For discovery, three nitrofen and three control fetal left lungs per time point were profiled on *Arraystar Rat Circular RNA Microarrays* after RNase R enrichment. Raw data were processed with Agilent Feature Extraction and R, and differential expression was assessed with a linear model and Benjamini–Hochberg correction (adjusted $p < 0.05$). Back-splice junction coordinates for differentially expressed (DE) probes were mapped to the rat genome using *FindBacksplice*.[1] Using these genomic coordinates, we generated predicted mature circRNA sequences with a modified CRAFT pipeline, identified miRNA binding sites (miRanda; score > 140, energy < −20), and compiled candidate mRNA targets via multiMiR.[2] Functional enrichment of DE parental genes and predicted circRNA::miRNA::mRNA targets employed GO and KEGG. We corroborated these results using Oxford Nanopore RNA sequencing. After preprocessing, we tested differential expression with two-sided t-tests and Benjamini–Hochberg correction, then compared observed DE genes to predicted circRNA::miRNA::mRNA targets and repeated GO/KEGG analyses to confirm pathway-level concordance.

For molecular validation and quantification, total RNA and genomic DNA were isolated with on-column DNase/RNase steps. We confirmed circRNA circularity by divergent primer PCR across BSJs on RNase R–treated cDNA, pJET subcloning, and Sanger sequencing (directional PCR to verify internal exon order where applicable). RT-qPCR quantified circRNAs and their parental mRNAs from the same cDNA pool. Spatial localization was assessed by BaseScope™ in situ hybridization using target-specific probes.

Finally, we performed a cross-species comparison by aligning rat circRNA findings (including the circAnp32e mature sequence) and enriched pathways with previously published human CDH circRNA microarray datasets at mid- and end-gestation[3], highlighting conserved sequence features and biology. Experimental details and analysis code are provided in the following section, as well as in Tables E1–E5.

*Circular RNA Microarray*

RNA was extracted (RNeasy Kit, Ref. 74104, Qiagen), quantified and quality assessed (NanoDrop ND-1000, Thermo Scientific), and checked for integrity by denaturing agarose gel electrophoresis. CircRNAs were enriched by RNase R digestion (Epicentre, Inc.), labeled using random primers (Arraystar Super RNA Labeling Protocol), hybridized to Arraystar Rat Circular RNA Microarrays (Ref. AS-S-CR-R-V2.0, Arraystar Inc.), incubated at 65°C for 17h, and scanned on an Agilent Scanner G2505C. Data were processed using Agilent Feature Extraction Software (v11.0.1.1) and R. Detection thresholds were determined via background signal slope analysis.

*Bioinformatics analysis and modified CRAFT*

Differential expression analysis of the circRNA microarray expression data was performed using a linear model as implemented by the *lm()* function, and Benjamini-Hochberg multiple testing correction performed with *p.adjust()*, both in the R package, *stats*, version 4.4.1. An adjusted p-value < 0.05 was used for determining differential expression. Observations with incomplete data were excluded from analysis.

Back-splice coordinates of DE circRNAs were generated using our published algorithm *FindBacksplice*.[1] To provide downstream circRNA::miRNA::mRNA target prediction, the CRAFT pipeline was modified to fit system requirements and tool availability. Putative circRNA mature sequences were produced from the CRAFT script *script_sequence_extraction.sh*. MicroRNA binding sites were predicted with miRanda v3.3 (pairing score >140, energy <−20) as implemented in the CRAFT script *script_miRNA_detection.sh*. Downstream targets were identified with the R library multiMiR version 1.26.0.

Finally, we performed GO and KEGG pathway analyses of DE parental genes and circRNA::miRNA::mRNA targets as well as on the Oxford Nanopore RNA sequencing datasets. Pathway analysis was carried out in R v4.4.1 with the libraries KEGGREST v1.44.1 and GO.db v3.19.1, representing a snapshot of the Gene Ontology on January 17, 2024.

The corresponding code available on our public GitHub repository at [https://github.com/m-kraljevic/modified-CRAFT-circRNA-CDH-pipeline](https://github.com/m-kraljevic/modified-CRAFT-circRNA-CDH-pipeline)

*RNA isolation, RNase R treatment, and DNA isolation*

RNA was extracted from fetal lungs at E15 and E21 using PureLink™ RNA Mini Kit (Ref. 12183018A, Invitrogen) by on-column DNase I treatment (Ref. 12185010, Invitrogen). RNA was quantified (NanoDrop ND-1000) and visualized on a 2% bleach agarose gel. To enrich for circRNAs, 2–4 µg RNA was digested with RNase R (25U; Ref. RNR07250, Lucigen) by incubating at 65°C (5–10 min), 37°C (20 min), and 70°C (10 min). Genomic DNA was isolated using PureLink™ Genomic DNA Mini Kit (Ref. K1820-01, Invitrogen) with in-column RNase A digestion.

*cDNA library generation, circRNA backsplice-junction detection, subcloning, and Sanger sequencing*

cDNA synthesis was performed with Maxima H Minus Reverse Transcriptase (Ref. EP0751, Thermo Scientific). Primers were designed via Primer3Web (version 4.1.0; Table E2). PCR was carried out with 2X DreamTaq PCR Master Mix (Ref. K1071, Thermo Scientific™) on a Veriti™ 96-Well Thermal Cycler (Applied Biosystems). Directional PCRs used Q5 High-Fidelity DNA Polymerase (Ref. M0491S, NEB) on 1:10 or 1:20 diluted RNase R-treated cDNA. PCR products were separated on 2.5% agarose gels, stained with SYBR™ Gold Nucleic Acid Gel Stain (Ref. E11494, Invitrogen), purified (PureLink™ Quick Gel Extraction Kit, Ref. K210012, Invitrogen), and subcloned with CloneJET PCR Cloning Kit (Ref. K1232, Thermo Scientific™). Ligation products were transformed into NEB 5-alpha Competent E. coli (Ref. C2988J, NEB) and selected on LB-ampicillin plates. Positive colonies were expanded, after which plasmids were extracted (GeneJET Plasmid Miniprep Kit, Ref. K0502, Thermo Scientific™) and sequenced (TCAG, SickKids, Toronto). Sequences were analyzed with ApE, plasmid editor [8] and 4Peaks software (Nucleobytes B.V., Aalsmeer, The Netherlands).

*RT-qPCR*

Following primer optimization and sequence verification, RT-qPCR was performed with SsoAdvanced Universal SYBR Green Supermix (Ref. 1725271, BioRad). cDNA was diluted 1:100 for parental genes and 1:10–1:20 for circRNAs. Expression was normalized to GAPDH using the ΔΔCt method. Details are provided in Table E2.

*Oxford Nanopore sequencing*

RNA libraries were prepared using the direct RNA sequencing kit (SQK-RNA004) (Oxford Nanopore Technologies) following the recommended protocol. A PromethION 24 device was used for sequencing with Flow Cells (FLO-PRO004RA). The flow cells were thawed at room temperature for 20 minutes, and available pore numbers were checked prior to loading the libraries. For detailed protocol of the Oxford Nanopore sequencing platform see Table E5. A two-sided t-test was performed in Python for each of the time points on the Oxford Nanopore RNA sequencing data. After applying Benjamini-Hochberg multiple testing correction, E21 had 1018 DE genes.

1. Kraljevic M, Ozturk A, Jank M, LeDuc RD, Keijzer R. FindBacksplice: a Tool for Locating Circular RNA Backsplice Coordinates [Internet]. bioRxiv; 2025 [cited 2025 Sept 15]. p. 2025.09.08.674962. Available from: https://www.biorxiv.org/content/10.1101/2025.09.08.674962v1

2. Dal Molin A, Gaffo E, Difilippo V, Buratin A, Tretti Parenzan C, Bresolin S, et al. CRAFT: a bioinformatics software for custom prediction of circular RNA functions. Brief Bioinform. 2022 Mar 10;23(2):bbab601.

3. Wagner R, Jha A, Ayoub L, Kahnamoui S, Patel D, Mahood TH, et al. Can circular RNAs be used as prenatal biomarkers for congenital diaphragmatic hernia? European Respiratory Journal [Internet]. 2020 Feb 1 [cited 2022 July 21];55(2). Available from: https://erj.ersjournals.com/content/55/2/1900514

#### Supplementary Figures

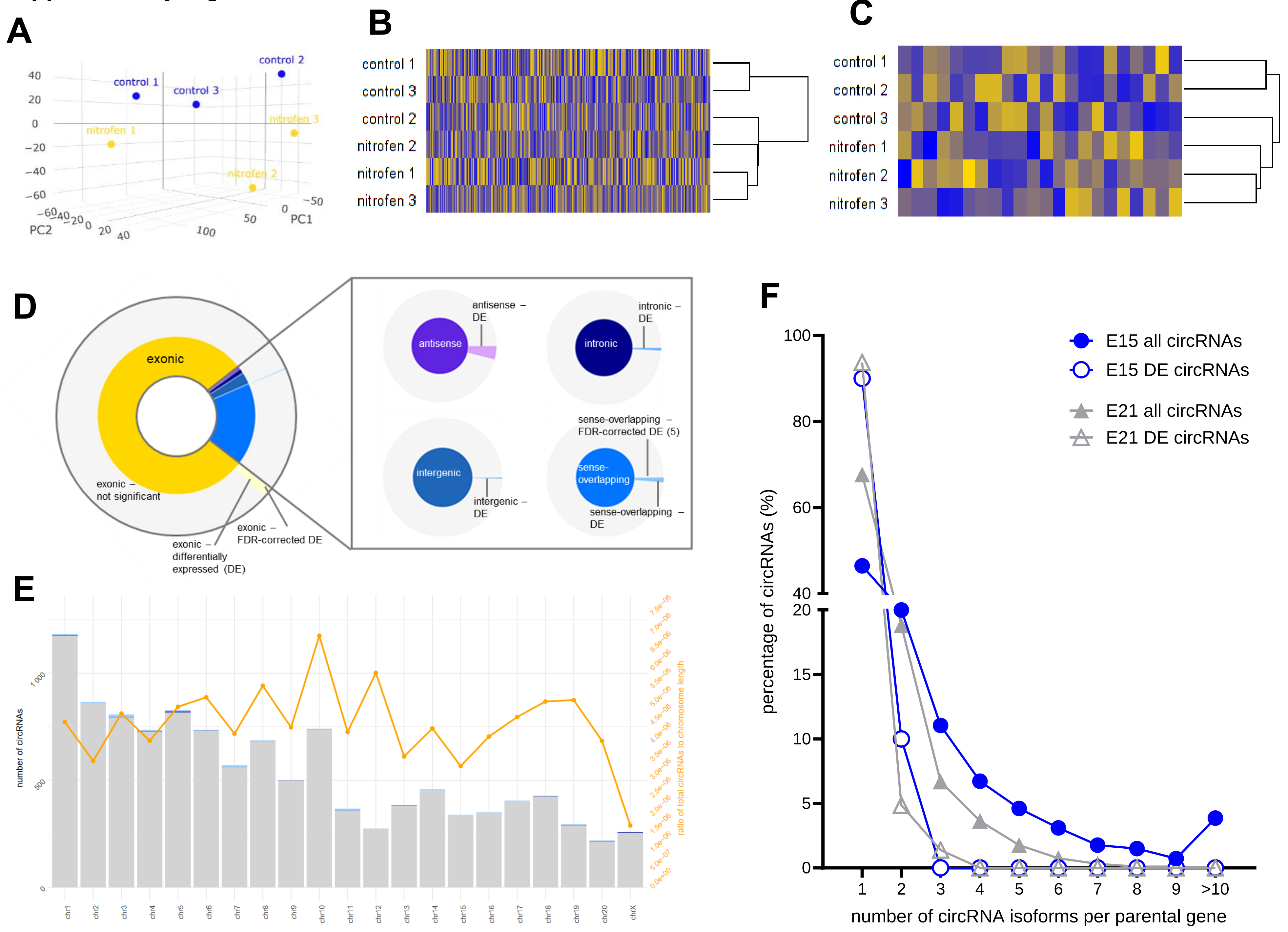

**Supplementary Figure E1 – Unique biosignature for lung hypoplasia in CDH**

Control and nitrofen-induced lung samples at embryonic day (E)15 segregate based on their overall circRNA profile plotted on a principal component analysis (A). The heatmap reveals that CDH lung datasets cluster closely together indicating a similar circRNA profile. Among the controls, control lung 2 is more similar to the nitrofen group in its total and differentially expressed (DE) circRNA profile while controls 1 and 3 exhibit distinct clustering (B and C). Distribution of DE circRNAs at E15 resembles late gestation with the majority being exonic (77%) or sense overlapping (23%) (D). Numbers of circRNAs across all chromosomes and the ratio of circRNAs to the chromosome length reveal a similar pattern to E21 (see Figure 1G), with the highest circRNA enrichment on chromosomes 10, 8, and 12 (E). CircRNAs per parental gene at early and late gestation show a higher number of total circRNA isoforms per parental gene at E15 (F).

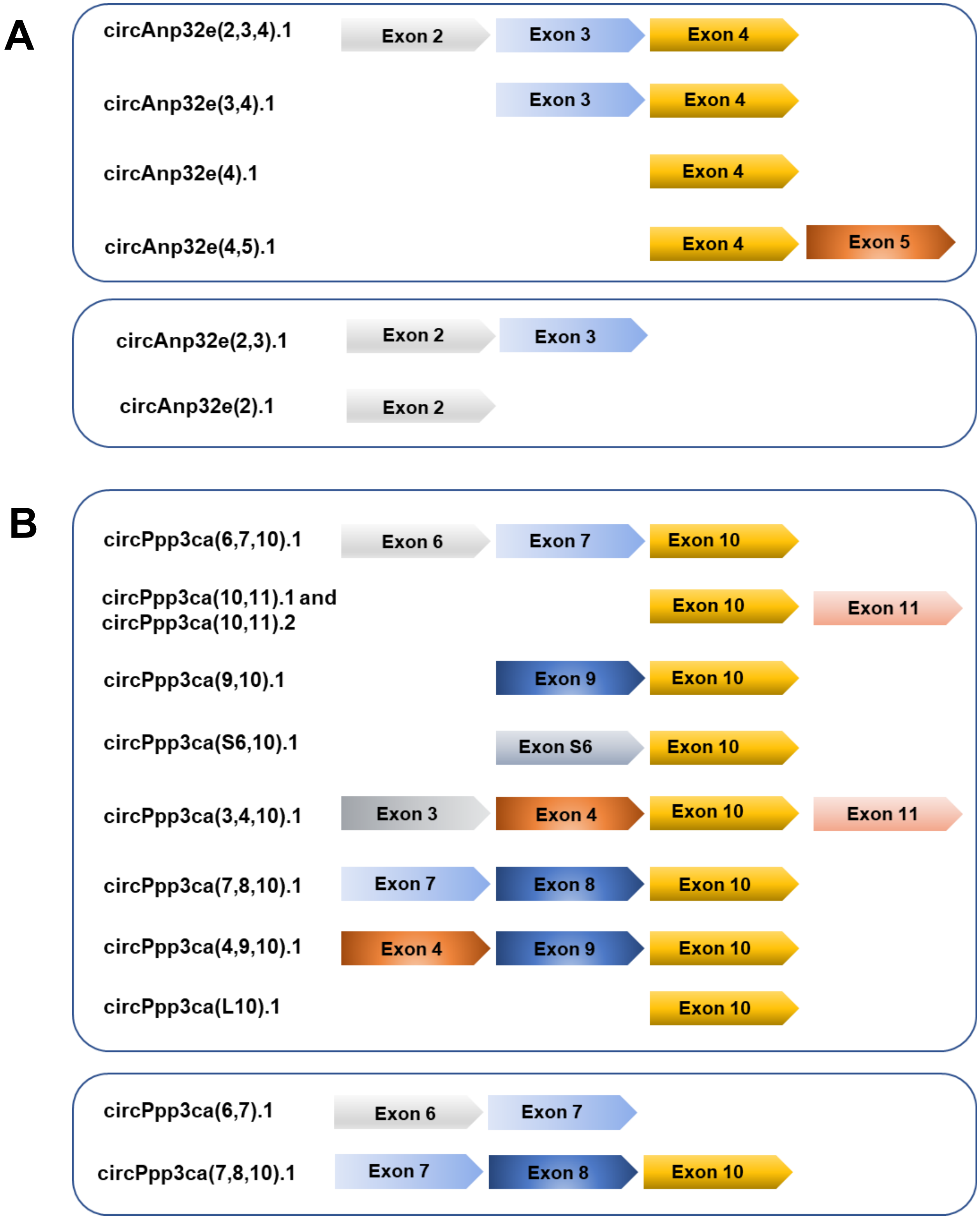


**Supplementary Figure E2 – Multiple circRNA isoforms derived from the parental genes *anp32e* and *ppp3ca* curated for the species rat by *circAtlas***

Investigation of multiple circRNA isoforms derived from the parental genes *anp32e* (A) and *ppp3ca* (B) curated for the rat species in the circAtlas database (accessed 11.11.2024). Only isoforms with back-splice junctions involving the same exons as our prime candidates/isoforms are shown.

A

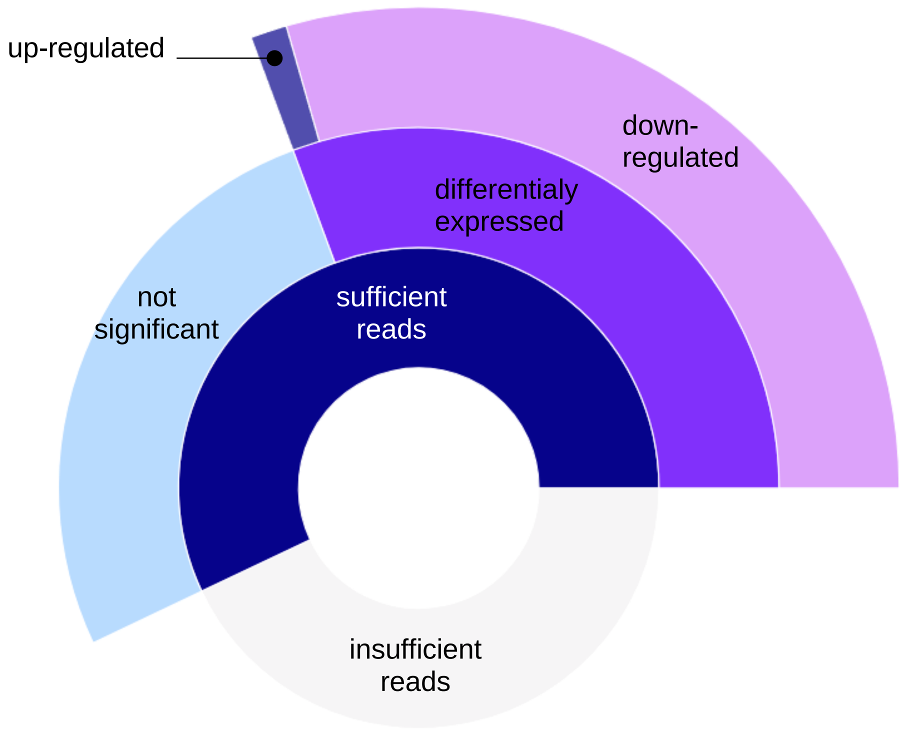


B

| gene | RNAseq | log fold change | p-value | adjusted p-value |
|---|---|---|---|---|
| *anp32e* | n.s. | -3.05 | 0.08 | 0.15 |
| *tial1* | down | -3.74 | 0.001 | 0.016 |
| *larp4b* | down | -1.56 | 0.01 | 0.04 |

**Supplementary Figure E3 – Detection of linear host genes *anp32e*, *ppp3ca* and *larp4b***

RNA sequencing was used to assess expression of parental genes of circRNAs: 57% of all parental genes at E21 were detected by Oxford Nanopore sequencing and 30% were DE (A), including *Tial1* confirming our previously published qPCR data. *Anp32e* appeared to be downregulated, but did not reach statistical significance likely due to small sample size (N=3 control and N=3 nitrofen-induced CDH lungs) (B).

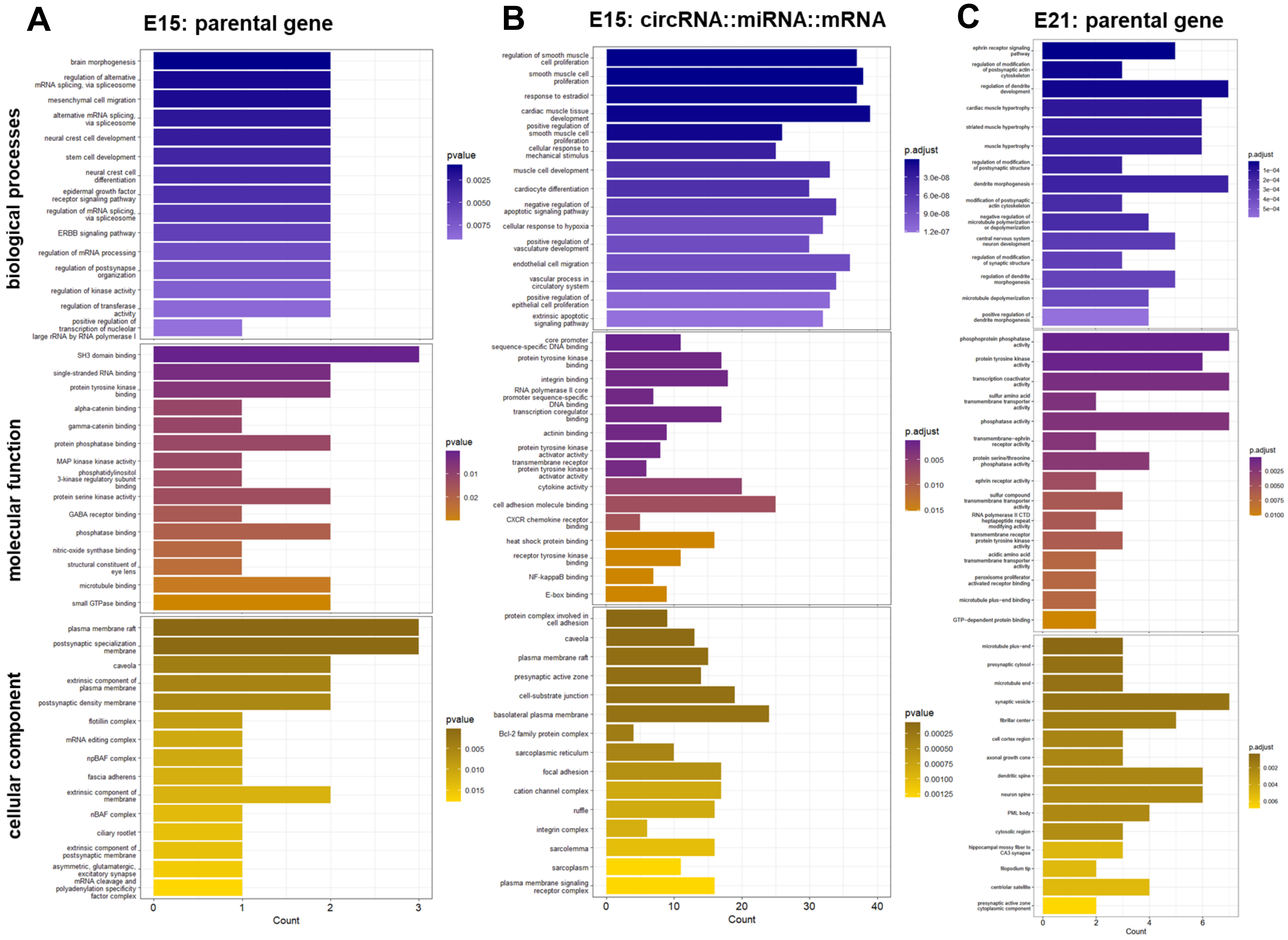

A
E15: parental gene
biological processes
brain morphogenesis
regulation of alternative mRNA splicing, via spliceosome
mesenchymal cell migration
alternative mRNA splicing, via spliceosome
neural crest cell development
stem cell development
neural crest cell differentiation
epidermal growth factor receptor signaling pathway
regulation of mRNA splicing, via spliceosome
ERBB signaling pathway
regulation of mRNA processing
regulation of postsynapse organization
regulation of kinase activity
regulation of transferase activity
positive regulation of transcription of nucleolar large rRNA by RNA polymerase I
pvalue
0.0025
0.0050
0.0075
molecular function
SH3 domain binding
single-stranded RNA binding
protein tyrosine kinase binding
alpha-catenin binding
gamma-catenin binding
protein phosphatase binding
MAP kinase kinase activity
phosphatidylinositol 3-kinase regulatory subunit binding
protein serine kinase activity
GABA receptor binding
phosphatase binding
nitric-oxide synthase binding
structural constituent of eye lens
microtubule binding
small GTPase binding
pvalue
0.01
0.02
cellular component
plasma membrane raft
postsynaptic specialization membrane
caveola
extrinsic component of plasma membrane
postsynaptic density membrane
flotillin complex
mRNA editing complex
npBAF complex
fascia adherens
extrinsic component of membrane
nBAF complex
ciliary rootlet
extrinsic component of postsynaptic membrane
asymmetric, glutamatergic, excitatory synapse
mRNA cleavage and polyadenylation specificity factor complex
pvalue
0.005
0.010
0.015
0
1
2
3
Count
B
E15: circRNA::miRNA::mRNA
regulation of smooth muscle cell proliferation
smooth muscle cell proliferation
response to estradiol
cardiac muscle tissue development
positive regulation of smooth muscle cell proliferation
cellular response to mechanical stimulus
muscle cell development
cardiocyte differentiation
negative regulation of apoptotic signaling pathway
cellular response to hypoxia
positive regulation of vasculature development
endothelial cell migration
vascular process in circulatory system
positive regulation of epithelial cell proliferation
extrinsic apoptotic signaling pathway
p.adjust
3.0e-08
6.0e-08
9.0e-08
1.2e-07
core promoter sequence-specific DNA binding
protein tyrosine kinase binding
integrin binding
RNA polymerase II core promoter sequence-specific DNA binding
transcription coregulator binding
actinin binding
protein tyrosine kinase activator activity
transmembrane receptor protein tyrosine kinase activator activity
cytokine activity
cell adhesion molecule binding
CXCR chemokine receptor binding
heat shock protein binding
receptor tyrosine kinase binding
NF-kappaB binding
E-box binding
p.adjust
0.005
0.010
0.015
protein complex involved in cell adhesion
caveola
plasma membrane raft
presynaptic active zone
cell-substrate junction
basolateral plasma membrane
Bcl-2 family protein complex
sarcoplasmic reticulum
focal adhesion
cation channel complex
ruffle
integrin complex
sarcolemma
sarcoplasm
plasma membrane signaling receptor complex
pvalue
0.00025
0.00050
0.00075
0.00100
0.00125
0
10
20
30
40
Count
C
E21: parental gene
ephrin receptor signaling pathway
regulation of modification of postsynaptic actin cytoskeleton
regulation of dendrite development
cardiac muscle hypertrophy
striated muscle hypertrophy
muscle hypertrophy
regulation of modification of postsynaptic structure
dendrite morphogenesis
modification of postsynaptic actin cytoskeleton
negative regulation of microtubule polymerization or depolymerization
central nervous system neuron development
regulation of modification of synaptic structure
regulation of dendrite morphogenesis
microtubule depolymerization
positive regulation of dendrite morphogenesis
p.adjust
1e-04
2e-04
3e-04
4e-04
5e-04
phosphoprotein phosphatase activity
protein tyrosine kinase activity
transcription coactivator activity
sulfur amino acid transmembrane transporter activity
phosphatase activity
transmembrane-ephrin receptor activity
protein serine/threonine phosphatase activity
ephrin receptor activity
sulfur compound transmembrane transporter activity
RNA polymerase II CTD heptapeptide repeat modifying activity
transmembrane receptor protein tyrosine kinase activity
acidic amino acid transmembrane transporter activity
peroxisome proliferator activated receptor binding
microtubule plus-end binding
GTP-dependent protein binding
p.adjust
0.0025
0.0050
0.0075
0.0100
microtubule plus-end
presynaptic cytosol
microtubule end
synaptic vesicle
fibrillar center
cell cortex region
axonal growth cone
dendritic spine
neuron spine
PML body
cytosolic region
hippocampal mossy fiber to CA3 synapse
filopodium tip
centriolar satellite
presynaptic active zone cytoplasmic component
p.adjust
0.002
0.004
0.006
0
2
4
6
Count

**Supplementary Figure E4 – Enrichment for pathways related to hypoxia response, cell proliferation, vascular processes, and inflammation in circRNA::miRNA::mRNA downstream targets at E15**

The parental genes of DE circRNAs at E15 enriched pathways related to splicing/mRNA processing, and (mesenchymal, stem cell and neural crest) cell development as well as migration. Augmented molecular functions mainly involved binding activity, while impacted cellular components comprised membrane structures as well as complexes regulating gene expression and RNA processing (A). Next, we investigated potential biological function of circRNAs in early stages of abnormal lung development by circRNA::miRNA::mRNA interaction. The 730 target genes displayed increased enrichment for (smooth muscle, epithelial and endothelial) proliferation and migration, response to hypoxia, and vascular development (B). At E21, parental genes were associated with GO terms related to muscle hypertrophy and neural development/synapses (C).

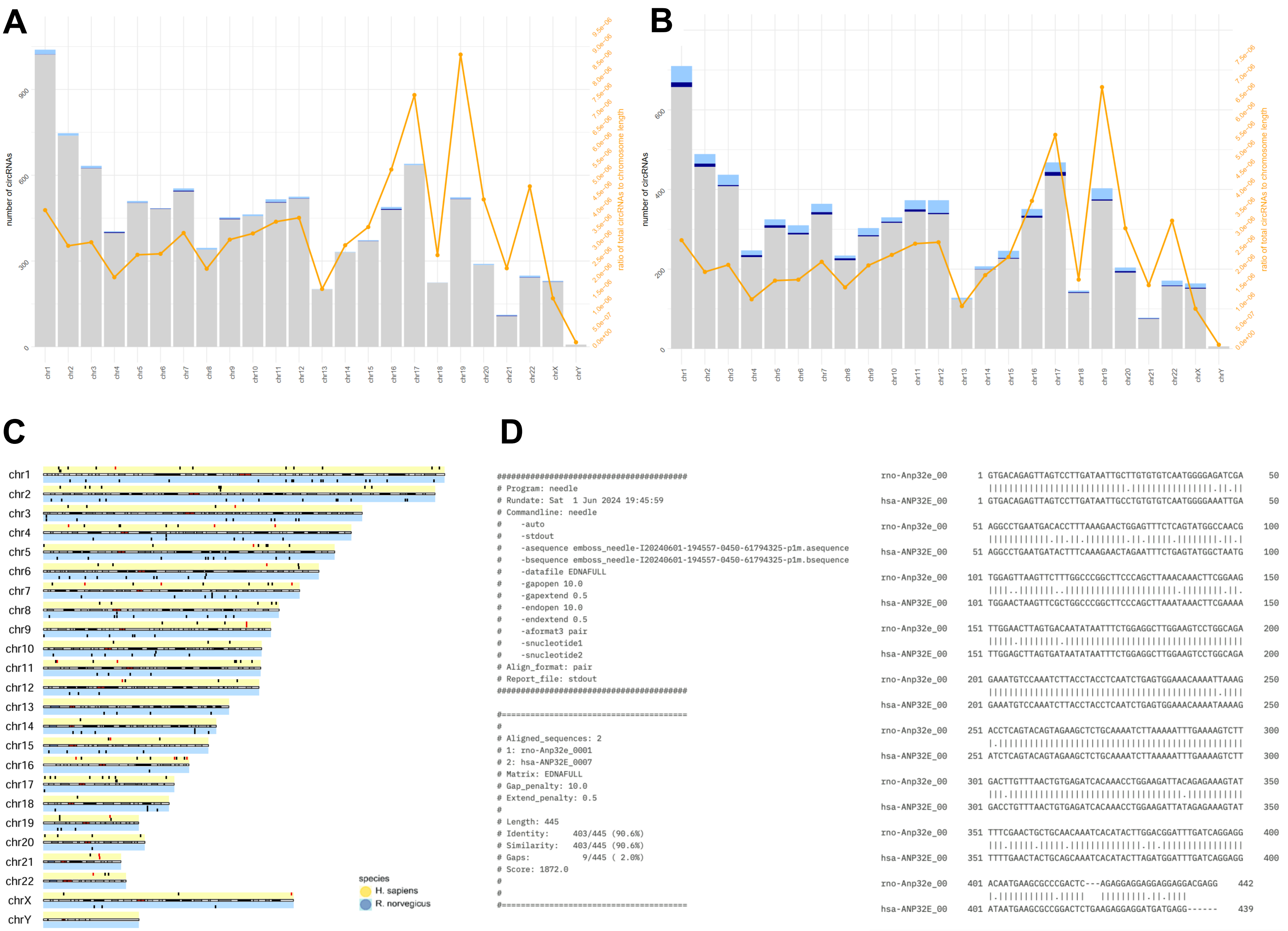
A
number of circRNAs
ratio of total circRNAs to chromosome length
0
300
600
900
0.0e+00
5.0e-07
1.0e-06
1.5e-06
2.0e-06
2.5e-06
3.0e-06
3.5e-06
4.0e-06
4.5e-06
5.0e-06
5.5e-06
6.0e-06
6.5e-06
7.0e-06
7.5e-06
8.0e-06
8.5e-06
9.0e-06
9.5e-06
chr1 chr2 chr3 chr4 chr5 chr6 chr7 chr8 chr9 chr10 chr11 chr12 chr13 chr14 chr15 chr16 chr17 chr18 chr19 chr20 chr21 chr22 chrX chrY
B
number of circRNAs
ratio of total circRNAs to chromosome length
0
200
400
600
0.0e+00
5.0e-07
1.0e-06
1.5e-06
2.0e-06
2.5e-06
3.0e-06
3.5e-06
4.0e-06
4.5e-06
5.0e-06
5.5e-06
6.0e-06
6.5e-06
7.0e-06
7.5e-06
chr1 chr2 chr3 chr4 chr5 chr6 chr7 chr8 chr9 chr10 chr11 chr12 chr13 chr14 chr15 chr16 chr17 chr18 chr19 chr20 chr21 chr22 chrX chrY
C
chr1
chr2
chr3
chr4
chr5
chr6
chr7
chr8
chr9
chr10
chr11
chr12
chr13
chr14
chr15
chr16
chr17
chr18
chr19
chr20
chr21
chr22
chrX
chrY
species
H. sapiens
R. norvegicus
D
########################################
### Program: needle
### Rundate: Sat  1 Jun 2024 19:45:59
### Commandline: needle
#    -auto
#    -stdout
#    -asequence emboss_needle-I20240601-194557-0450-61794325-p1m.asequence
#    -bsequence emboss_needle-I20240601-194557-0450-61794325-p1m.bsequence
#    -datafile EDNAFULL
#    -gapopen 10.0
#    -gapextend 0.5
#    -endopen 10.0
#    -endextend 0.5
#    -aformat3 pair
#    -snucleotide1
#    -snucleotide2
### Align_format: pair
### Report_file: stdout
########################################
#=======================================
#
### Aligned_sequences: 2
### 1: rno-Anp32e_0001
### 2: hsa-ANP32E_0007
### Matrix: EDNAFULL
### Gap_penalty: 10.0
### Extend_penalty: 0.5
#
### Length: 445
# Identity:     403/445 (90.6%)
# Similarity:   403/445 (90.6%)
# Gaps:           9/445 ( 2.0%)
### Score: 1872.0
#
#
#=======================================
rno-Anp32e_00      1 GTGACAGAGTTAGTCCTTGATAATTGCTTGTGTGTCAATGGGGAGATCGA     50
hsa-ANP32E_00      1 GTGACAGAGTTAGTCCTTGATAATTGCCTGTGTGTCAATGGGGAAATTGA     50
rno-Anp32e_00     51 AGGCCTGAATGACACCTTTAAAGAACTGGAGTTTCTCAGTATGGCCAACG    100
hsa-ANP32E_00     51 AGGCCTGAATGATACTTTCAAAGAACTAGAATTTCTGAGTATGGCTAATG    100
rno-Anp32e_00    101 TGGAGTTAAGTTCTTTGGCCCGGCTTCCCAGCTTAAACAAACTTCGGAAG    150
hsa-ANP32E_00    101 TGGAACTAAGTTCGCTGGCCCGGCTTCCCAGCTTAAATAAACTTCGAAAA    150
rno-Anp32e_00    151 TTGGAACTTAGTGACAATATAATTTCTGGAGGCTTGGAAGTCCTGGCAGA    200
hsa-ANP32E_00    151 TTGGAGCTTAGTGATAATATAATTTCTGGAGGCTTGGAAGTCCTGGCAGA    200
rno-Anp32e_00    201 GAAATGTCCAAATCTTACCTACCTCAATCTGAGTGGAAACAAAATTAAAG    250
hsa-ANP32E_00    201 GAAATGTCCAAATCTTACCTACCTCAATCTGAGTGGAAACAAAATAAAAG    250
rno-Anp32e_00    251 ACCTCAGTACAGTAGAAGCTCTGCAAAATCTTAAAAATTTGAAAAGTCTT    300
hsa-ANP32E_00    251 ATCTCAGTACAGTAGAAGCTCTGCAAAATCTTAAAAATTTGAAAAGTCTT    300
rno-Anp32e_00    301 GACTTGTTTAACTGTGAGATCACAAACCTGGAAGATTACAGAGAAAGTAT    350
hsa-ANP32E_00    301 GACCTGTTTAACTGTGAGATCACAAACCTGGAAGATTATAGAGAAAGTAT    350
rno-Anp32e_00    351 TTTCGAACTGCTGCAACAAATCACATACTTGGACGGATTTGATCAGGAGG    400
hsa-ANP32E_00    351 TTTTGAACTACTGCAGCAAATCACATACTTAGATGGATTTGATCAGGAGG    400
rno-Anp32e_00    401 ACAATGAAGCGCCCGACTC---AGAGGAGGAGGAGGAGGACGAGG    442
hsa-ANP32E_00    401 ATAATGAAGCGCCGGACTCTGAAGAGGAGGATGATGAGG------    439

**Supplementary Figure E5 – circRNAs in human CDH and highly conserved mature circAnp32e sequence across species**

The ratio of total circRNA number to the chromosome length at mid- (A) and end-gestation (B) showed the highest enrichment for chromosomes 17 and 19. Chromosomal locations of differentially expressed (DE) circRNA in rat and human CDH were plotted to detect potential "hot spots" of splice sites resulting in circRNA formation with red being mid-gestation/E15 and black end-gestation/E21 (C). Alignment of the circAnp32e mature sequence between the rat and human orthologues indicated a 90% sequence identity (D).

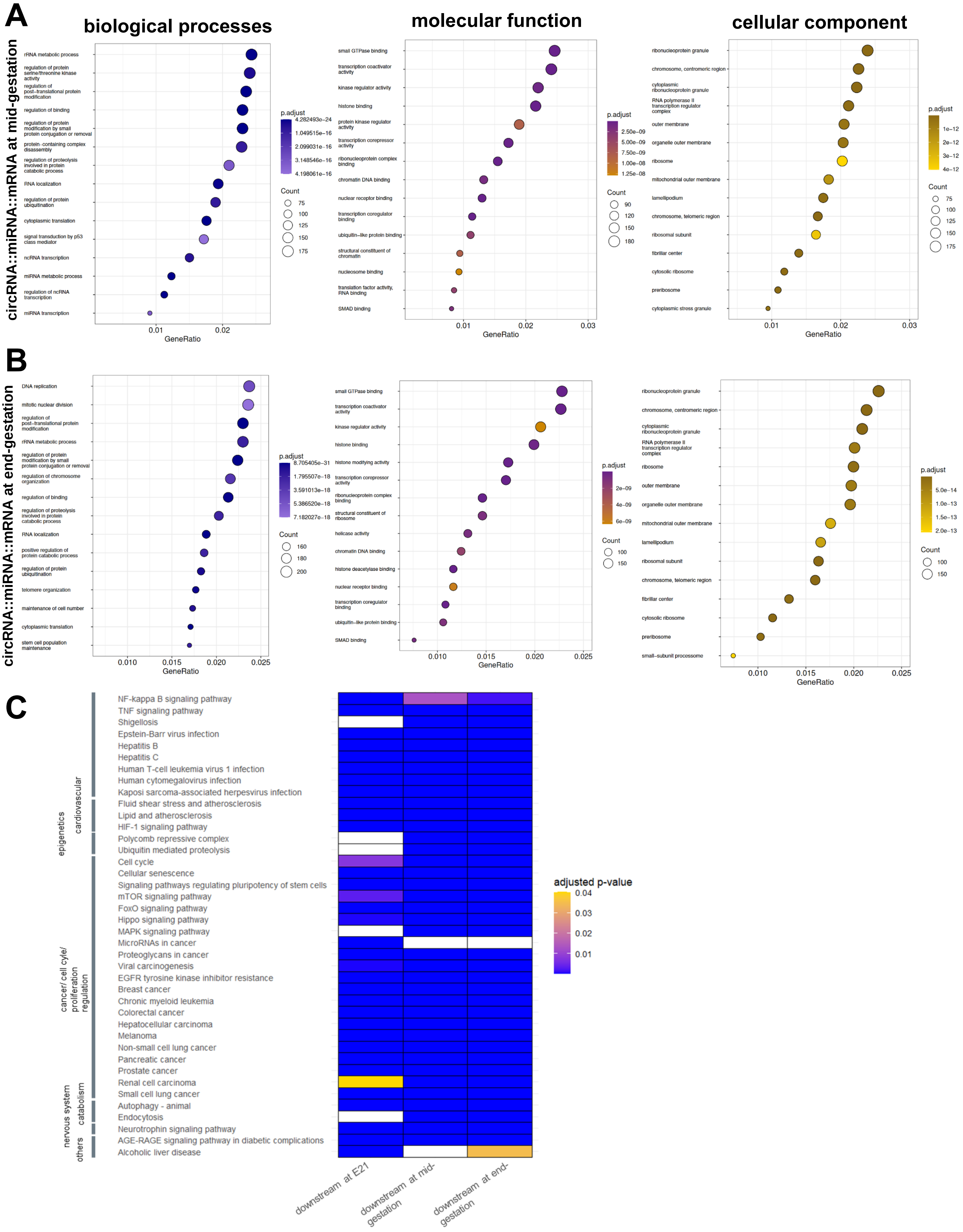
A
biological processes
molecular function
cellular component
circRNA::miRNA::mRNA at mid-gestation
rRNA metabolic process
regulation of protein serine/threonine kinase activity
regulation of post−translational protein modification
regulation of binding
regulation of protein modification by small protein conjugation or removal
protein−containing complex disassembly
regulation of proteolysis involved in protein catabolic process
RNA localization
regulation of protein ubiquitination
cytoplasmic translation
signal transduction by p53 class mediator
ncRNA transcription
miRNA metabolic process
regulation of ncRNA transcription
miRNA transcription
GeneRatio
p.adjust
4.282493e−24
1.049515e−16
2.099031e−16
3.148546e−16
4.198061e−16
Count
75
100
125
150
175
small GTPase binding
transcription coactivator activity
kinase regulator activity
histone binding
protein kinase regulator activity
transcription corepressor activity
ribonucleoprotein complex binding
chromatin DNA binding
nuclear receptor binding
transcription coregulator binding
ubiquitin−like protein binding
structural constituent of chromatin
nucleosome binding
translation factor activity, RNA binding
SMAD binding
2.50e−09
5.00e−09
7.50e−09
1.00e−08
1.25e−08
90
120
150
180
ribonucleoprotein granule
chromosome, centromeric region
cytoplasmic ribonucleoprotein granule
RNA polymerase II transcription regulator complex
outer membrane
organelle outer membrane
ribosome
mitochondrial outer membrane
lamellipodium
chromosome, telomeric region
ribosomal subunit
fibrillar center
cytosolic ribosome
preribosome
cytoplasmic stress granule
1e−12
2e−12
3e−12
4e−12
B
circRNA::miRNA::mRNA at end-gestation
DNA replication
mitotic nuclear division
regulation of post−translational protein modification
rRNA metabolic process
regulation of protein modification by small protein conjugation or removal
regulation of chromosome organization
regulation of binding
regulation of proteolysis involved in protein catabolic process
RNA localization
positive regulation of protein catabolic process
regulation of protein ubiquitination
telomere organization
maintenance of cell number
cytoplasmic translation
stem cell population maintenance
8.705405e−31
1.795507e−18
3.591013e−18
5.386520e−18
7.182027e−18
160
180
200
histone modifying activity
helicase activity
histone deacetylase binding
2e−09
4e−09
6e−09
100
150
small−subunit processome
5.0e−14
1.0e−13
1.5e−13
2.0e−13
C
cardiovascular
epigenetics
cancer/ cell cyle/ proliferation regulation
catabolism
nervous system
others
NF-kappa B signaling pathway
TNF signaling pathway
Shigellosis
Epstein-Barr virus infection
Hepatitis B
Hepatitis C
Human T-cell leukemia virus 1 infection
Human cytomegalovirus infection
Kaposi sarcoma-associated herpesvirus infection
Fluid shear stress and atherosclerosis
Lipid and atherosclerosis
HIF-1 signaling pathway
Polycomb repressive complex
Ubiquitin mediated proteolysis
Cell cycle
Cellular senescence
Signaling pathways regulating pluripotency of stem cells
mTOR signaling pathway
FoxO signaling pathway
Hippo signaling pathway
MAPK signaling pathway
MicroRNAs in cancer
Proteoglycans in cancer
Viral carcinogenesis
EGFR tyrosine kinase inhibitor resistance
Breast cancer
Chronic myeloid leukemia
Colorectal cancer
Hepatocellular carcinoma
Melanoma
Non-small cell lung cancer
Pancreatic cancer
Prostate cancer
Renal cell carcinoma
Small cell lung cancer
Autophagy - animal
Endocytosis
Neurotrophin signaling pathway
AGE-RAGE signaling pathway in diabetic complications
Alcoholic liver disease
adjusted p-value
0.04
0.03
0.02
0.01
downstream at E21
downstream at mid-gestation
downstream at end-gestation

**Supplementary Figure E6 – Strong overlap in predicted biological function (miRNA sponging) of circRNAs between rat E21 and human CDH at mid- and end-gestation.**

Predicted circRNA::miRNA::mRNA at both mid- (A) and end-gestation (B) showed enrichment for biological processes such as RNA regulation, epigenetic pathways, and protein binding/modification in CDH lungs. In addition, KEGG pathway analyses revealed that downstream targets were associated with inflammation and infection (*e.g.*, NF-κB and TNF signaling as well as viral infections), regulation of cell cycle and proliferation, and cardiovascular pathways. We observed a strong overlap in the top 15 enriched KEGG terms between rat E21 and human mid- and end-gestation CDH lungs (C).

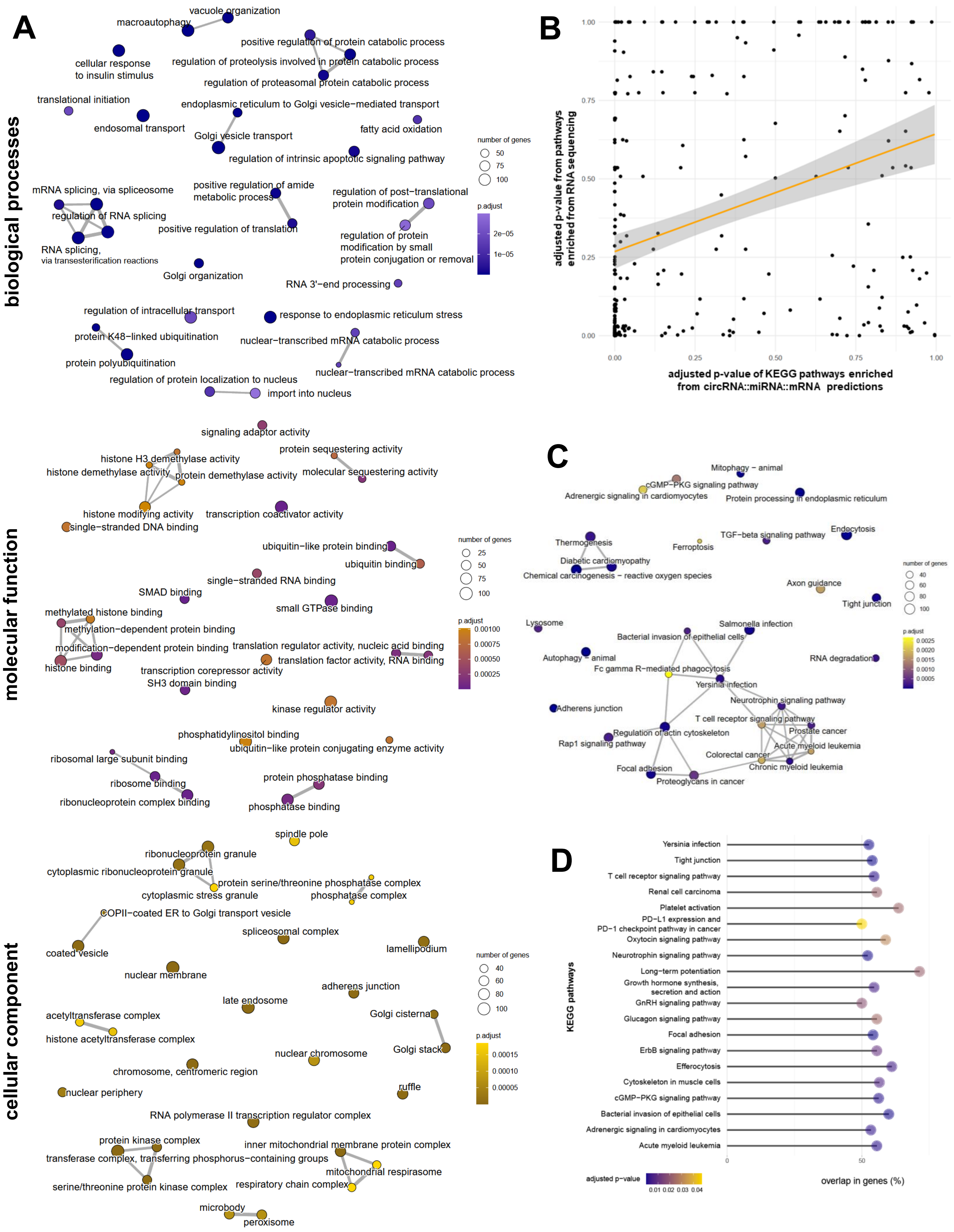
A
biological processes
vacuole organization
macroautophagy
positive regulation of protein catabolic process
cellular response
to insulin stimulus
regulation of proteolysis involved in protein catabolic process
regulation of proteasomal protein catabolic process
translational initiation
endoplasmic reticulum to Golgi vesicle−mediated transport
endosomal transport
fatty acid oxidation
Golgi vesicle transport
regulation of intrinsic apoptotic signaling pathway
mRNA splicing, via spliceosome
positive regulation of amide
metabolic process
regulation of post−translational
protein modification
regulation of RNA splicing
positive regulation of translation
regulation of protein
modification by small
protein conjugation or removal
RNA splicing,
via transesterification reactions
Golgi organization
RNA 3'−end processing
regulation of intracellular transport
response to endoplasmic reticulum stress
protein K48−linked ubiquitination
nuclear−transcribed mRNA catabolic process
protein polyubiquitination
nuclear−transcribed mRNA catabolic process
regulation of protein localization to nucleus
import into nucleus
number of genes
50
75
100
p.adjust
2e−05
1e−05
molecular function
signaling adaptor activity
protein sequestering activity
histone H3 demethylase activity
histone demethylase activity
protein demethylase activity
molecular sequestering activity
histone modifying activity
transcription coactivator activity
single−stranded DNA binding
ubiquitin−like protein binding
ubiquitin binding
single−stranded RNA binding
SMAD binding
small GTPase binding
methylated histone binding
methylation−dependent protein binding
modification−dependent protein binding
translation regulator activity, nucleic acid binding
translation factor activity, RNA binding
histone binding
transcription corepressor activity
SH3 domain binding
kinase regulator activity
phosphatidylinositol binding
ubiquitin−like protein conjugating enzyme activity
ribosomal large subunit binding
ribosome binding
protein phosphatase binding
ribonucleoprotein complex binding
phosphatase binding
number of genes
25
50
75
100
p.adjust
0.00100
0.00075
0.00050
0.00025
cellular component
spindle pole
ribonucleoprotein granule
cytoplasmic ribonucleoprotein granule
protein serine/threonine phosphatase complex
cytoplasmic stress granule
phosphatase complex
COPII−coated ER to Golgi transport vesicle
spliceosomal complex
coated vesicle
lamellipodium
nuclear membrane
adherens junction
late endosome
acetyltransferase complex
Golgi cisterna
histone acetyltransferase complex
nuclear chromosome
Golgi stack
chromosome, centromeric region
nuclear periphery
ruffle
RNA polymerase II transcription regulator complex
protein kinase complex
inner mitochondrial membrane protein complex
transferase complex, transferring phosphorus−containing groups
mitochondrial respirasome
serine/threonine protein kinase complex
respiratory chain complex
microbody
peroxisome
number of genes
40
60
80
100
p.adjust
0.00015
0.00010
0.00005
B
adjusted p-value from pathways
enriched from RNA sequencing
1.00
0.75
0.50
0.25
0.00
0.00
0.25
0.50
0.75
1.00
adjusted p-value of KEGG pathways enriched
from circRNA::miRNA::mRNA predictions
C
Mitophagy − animal
cGMP−PKG signaling pathway
Adrenergic signaling in cardiomyocytes
Protein processing in endoplasmic reticulum
Endocytosis
Thermogenesis
TGF−beta signaling pathway
Ferroptosis
Diabetic cardiomyopathy
Chemical carcinogenesis − reactive oxygen species
Axon guidance
Tight junction
Lysosome
Salmonella infection
Bacterial invasion of epithelial cells
Autophagy − animal
RNA degradation
Fc gamma R−mediated phagocytosis
Yersinia infection
Adherens junction
Neurotrophin signaling pathway
T cell receptor signaling pathway
Regulation of actin cytoskeleton
Prostate cancer
Rap1 signaling pathway
Acute myeloid leukemia
Colorectal cancer
Focal adhesion
Chronic myeloid leukemia
Proteoglycans in cancer
number of genes
40
60
80
100
p.adjust
0.0025
0.0020
0.0015
0.0010
0.0005
D
KEGG pathways
Yersinia infection
Tight junction
T cell receptor signaling pathway
Renal cell carcinoma
Platelet activation
PD−L1 expression and
PD−1 checkpoint pathway in cancer
Oxytocin signaling pathway
Neurotrophin signaling pathway
Long−term potentiation
Growth hormone synthesis,
secretion and action
GnRH signaling pathway
Glucagon signaling pathway
Focal adhesion
ErbB signaling pathway
Efferocytosis
Cytoskeleton in muscle cells
cGMP−PKG signaling pathway
Bacterial invasion of epithelial cells
Adrenergic signaling in cardiomyocytes
Acute myeloid leukemia
0
50
adjusted p−value
0.01 0.02 0.03 0.04
overlap in genes (%)

**Supplementary Figure E7 – Confirmation of predicted KEGG pathways using the modified CRAFT pipeline and Oxford Nanopore Sequencing**

Gene Ontology (GO) analyses of Oxford Nanopore Sequencing revealed networks related to post-transcriptional regulation including mRNA splicing, mRNA processing, and histone modification (A). p-values of all KEGG terms derived from the modified CRAFT pipeline were plotted against the pathways enriched by RNA sequencing, resulting in a moderate positive correlation (r=0.38; B). DE genes between nitrofen-induced and control lungs detected by Oxford Nanopore Sequencing showed enrichment for KEGG terms related to infection/immunity, cell adhesion (tight junction and adherence junction), and cancer (C). The proportion of predicted genes, confirmed to be DE based on RNA sequencing, was visualized for each pathway (D).

### Supplementary Tables

#### Supplementary Table E1 – Sex determination: primers and protocol

| **Primer/Protocol for sex determination** | |
|---|---|
| *Sry* forward | CCAGCATGCAGAATTCAGAGA |
| *Sry* reverse | CCCAGTCCTGTCCGTATATGAT |
| *Kdm5c* forward | AGACACCAGAGGCAGATAGA |
| *Kdm5c* reverse | CTCCATTAACAGGCGAGGAA |
| Protocol | Initial denaturation at 95°C for 2min;<br>Denaturation at 95°C for 30sec (35 cycles);<br>Annealing at 60°C for 30sec;<br>Extension at 72°C for 38sec;<br>Final extension at 72°C for 5min<br><br>The PCR products were analyzed by 1% agarose gel electrophoresis; the presence or absence of the *Sry* gene allowed for sex determination. |

**Supplementary Table E2 – Sequence validation: primers and protocols**

| **Primer** | **Sequence** | **Tm (°C)** | **product length** |
|---|---|---|---|
| *anp32e* forward | CAGGAGGACAATGAAGCGC | 58.89 | 158 bp |
| *anp32e* reverse | CCCTCTCCTACTTCTGAGCC | 58.59 | |
| *circAnp32e* forward | ACGGATTTGATCAGGAGGACA | 58.46 | 156 bp |
| *circAnp32e* reverse | GGCCATACTGAGAAACTCCAG | 59.12 | |
| *ppp3ca* forward | ACGTCCTGAACATCTGCTCA | 59.03 | 112 bp |
| *ppp3ca* reverse | CCTATTGCTCGGATCTTGTTCC | 58.87 | |
| *circPpp3ca* forward | TGAGATGCTGGTAAACGTCCT | 58.73 | 131 bp |
| *circPpp3ca* reverse | GCTGCAGGAAGTCACATACAG | 59.65 | |
| *Larp4b* forward | CAGAGCAGACAGACATGAGC | 59.08 | 104 bp |
| *Larp4b* reverse | CTGGCTTTCTGGTTGAGACTC | 59.25 | |
| *circLarp4b* forward | AGAATACGAACCTCTGCCTGA | 59.11 | 154 bp |
| *circLarp4b* reverse | CAGATGAGAGCTGTCCTTGC | 59.1 | |
| *IntAnp32e Ex 4* forward | TTGATCAGGAGGACAATGAAGC | 58.39 | 246 bp |
| *IntAnp32e Ex 3* reverse | GACTTCCAAGCCTCCAGAAATT | 58.57 | |
| *IntLarp4b Ex-5* forward | AGCAGACAGACATGAGCACT | 59.02 | 249 bp |
| *IntLarp4b Ex-3* reverse | GAGTCAATGCTGGGAGGGAA | 59.38 | |
| *GAPDH* forward | AAGAACCCTTCCGTAGCGA | 59.3 | 124 bp |
| *GAPDH* reverse | GCTCTCTGCTCCTCCCTGTTCTA | 58.9 | |

#### Supplementary Table E3 – Probes for BaseScope™ *in situ* hybridization

| Probe ID for circRNA detection and quantification | |
|---|---|
| circAnp32e | NPR-0043351 |
| circPpp3ca | NPR-0043352 |
| circLarp4b | NPR-0042589 |

#### Supplementary Table E4 – BaseScope protocol

| Step | Procedure |
|---|---|
| **Sectioning** | 5 µm tissue sections mounted on Superfrost™ Plus slides (Fisher Scientific) |
| **Baking (1)** | Slides baked at 60°C for 1 h |
| **Deparaffinization** | Xylene (2×5 min), 100% ethanol (2×2 min) |
| **Baking (2)** | Slides dried at 60°C for 5 min |
| **Peroxide Treatment** | Hydrogen peroxide (ACD, Ref. 322335), 10 min at room temperature |
| **Water Wash & Acclimation** | Washed twice in distilled water, then 10 sec in distilled water at 100°C |
| **Target Retrieval** | 15 min at 100°C in RNAscope™ Target Retrieval Solution (ACD, Ref. 322000) |
| **Rehydration** | Slides in distilled water (10 sec), 100% ethanol (3 min) |
| **Baking (3)** | Slides baked at 60°C for 5 min |
| **Barrier Application** | Hydrophobic barrier drawn around each section |
| **Protease Treatment** | Protease III (ACD, Ref. 321710), 15 min at 40°C in HybEZ™ II oven |
| **Wash (1)** | RNAscope™ Wash Buffer (ACD, Ref. 310091), 2×2 min |
| **Probe Hybridization** | BaseScope™ probes applied for 2 h at 40°C in HybEZ™ II oven |
| **Amplification Steps** | Amplifier 1–6: 15–30 min at 40°C; Amplifier 7: 60 min room temperature; Amplifier 8: 30 min room temperature; Wash buffer 2×2 min between each step |
| **Detection** | Fast RED for 20 min at room temperature in the dark |
| **Counterstaining** | Gill's hematoxylin (2 min), 0.02% ammonia water (2–3 dips) |
| **Baking (4)** | Slides dried at 60°C for 15 min |
| **Mounting** | VectaMount mounting medium (Vector Labs) |

**Supplementary Table E5 – Oxford Nanopore direct RNA library preparation and sequencing (SQK-RNA004)**

| Step | Details |
|---|---|
| **RTA ligation** | NEBNext® Quick Ligation Reaction Buffer (NEB B6058, 3 µL); RNaseOUT™ (Invitrogen 10777019, 1 µL); RTA (1 µL); T4 DNA Ligase (NEB M0202M, 1.5 µL); total RNA (1 µg). Incubate 10 min at room temperature. |
| **Add RT master mix** | To adaptor-ligated RNA add 23 µL master mix: nuclease-free water 9 µL; 10 mM dNTPs (NEB N0447) 2 µL; 5× First-Strand Buffer 8 µL; 0.1 M DTT 4 µL. Mix well. |
| **Add RT enzyme** | SuperScript™ III Reverse Transcriptase (Thermo Fisher 18080044), +2 µL per reaction; brief spin and mix. |
| **First-strand synthesis** | Thermocycler: 50 °C 50 min; 70 °C 10 min; hold 4 °C. |
| **Bead cleanup (post-RT)** | Mag-Bind® TotalPure NGS beads (Omega Bio-tek), 1.8× bead:sample; add 72 µL beads per RT mix; rotate 5 min; magnet 5 min (or until clear); wash 2× with 70% EtOH. |
| **Elute RNA–cDNA hybrids** | Elute in 23 µL nuclease-free water; resuspend pellet, rotate 5 min; elute on magnet. |
| **Sequencing adaptor ligation** | To 23 µL eluate add: NEBNext® Quick Ligation Reaction Buffer 8 µL; RNA Ligation Adapter (RLA) 6 µL; T4 DNA Ligase 3 µL. Incubate 10 min at RT. |
| **Final library cleanup & elution** | NGS beads 1× (40 µL); rotate 5 min; magnet 5 min; wash 2× with washing buffer; elute in 33 µL elution buffer. Use 1 µL for quant; 32 µL for sequencing (≤ 113 ng loaded). |
| **Device & flow cells** | PromethION 24; Flow Cells FLO-PRO004RA. Thaw at RT 20 min; check pore count; remove headspace air. |
| **Prime flow cells** | Load 500 µL priming mix (30 µL RNA flush tether + 1170 µL flow cell flush); after 5 min add another 500 µL. |
| **Prepare library mix & load** | Library mix 200 µL total: 32 µL final library + 100 µL Sequencing Buffer (SB) + 68 µL Library Solution (LIS). Load per ONT PromethION protocol. |
| **Sequencing** | Run up to 72 h; data collected as pod5 for downstream preprocessing. |

**Supplementary Table E6 – Strong overlap in the parental genes of circRNAs between rat and human CDH**

| | **Overlap between mid- and end-gestation** | **Overlap between mid-gestation and E21** | **Overlap between end-gestation and E21** | **Overlap between mid-, end-gestation and E21** |
|---|---|---|---|---|
| parental genes of all detected circRNAs | 4399 | 1055 | 883 | 882 |
| parental genes of DE circRNAs | 20 (*aff3, ambra1, anapc7, asap2, cntrob, col24a1, ddx17, dennd3, kcnh1, lmf1, lphn1, mkl1, pan3, phc3, pkd1l2, smyd3, tgfb1, trio, wdr37* and *zbtb40*) | 14 (*ambra1, bnip3l, dennd1a, erc1, eya3, nckap1, pan3, phc3, rapgef6, reep2, smyd3, spg7, susd1* and *zmym4*) | 31 (*abhd12, ambra1, arhgef12, armc8, cdc27, cdk14, clip2, cnot1, crim1, dtnb, eif4e3, ephb2, exoc4, hecw1, mapk1, pan3, pde4d, phc3, pkdcc, ppp3cc, ptpra, rtel1, scaf8, sec24a, slc4a7, smek2, smyd3, snx14, stk39, uso1* and *vps53)* | 4 (*ambra1, pan3, phc3* and *smyd3)* |
| parental gene function:<br>- Lung disease/ fibrosis/ SMCs | *mkl1, smyd3, tgfb1, ambra1* and *col24a1* | *bnip3l, smyd3,* and potentially *nckap1* | *ambra1, mapk1, pde4d, smyd3* and *ptpra* | *ambra1* and *smyd3* |
| - Embryonic development | *phc3* and *trio* | *eya3* and *phc3* | *ephb2* and *phc3* | *phc3* |
| - Cell proliferation: | *anapc7, cntrob, kcnh1, tgfb1* and *smyd3* | *ambra1, bnip3l, smyd3* and potentially *reep2* | *cdc27, cdk14, eif4e3, mapk1, pan3, ptpra* and *smyd3* | *ambra1, smyd3* and *pan3* |
| - RNA/DNA regulation: | *aff3, ddx17, pan3, phc3, smyd3* and *zbtb40* | *pan3, phc3, smyd3* and *zmym4* | *cnot1, eif4e3, pan3, phc3, smyd3, rtel1* and *scaf8* | *pan3, phc3* and *smyd3* |
| - Others: | *lphn1* (cellular adhesion) | | | |
